# Focal infrared stimulation modulates somatosensory cortical activity in mice: evidence for TRPV1 ion channel involvement

**DOI:** 10.1101/2025.09.15.676310

**Authors:** Zsófia Balogh-Lantos, Ágoston Csaba Horváth, Katalin Rozmer, Richárd Fiáth, Zsuzsanna Helyes, Zoltán Fekete

## Abstract

Infrared neural stimulation (INS) is a promising approach for minimally invasive modulation of neuronal activity, yet its cellular and molecular mechanisms in the intact brain remain incompletely understood. Here, we investigated focal INS in the somatosensory cortex of anesthetized mice using high-density Neuropixels recordings combined with histological analysis. Continuous-wave and pulsed (500 Hz) infrared illumination elicited robust modulation of cortical activity, with approximately half of recorded neurons exhibiting either increased or suppressed firing. The modulatory effects of stimulation were comparable across cortical layers and neuron types. While both stimulation modes were effective, continuous-wave stimulation typically produced stronger changes in firing rates and network dynamics than pulsed illumination. To explore molecular contributions, we compared responses in wild-type and TRPV1 knockout mice. Neuronal responses to INS were significantly reduced in knockout animals, indicating a key role for TRPV1 channels. We also observed layer- and neuron-type-specific differences in firing rate modulation between wild-type and knockout animals. Histological analysis confirmed that TRPV1-expressing neurons are distributed throughout cortical layers, supporting their involvement in the observed responses. These findings provide direct evidence that TRPV1 contributes to INS-evoked cortical activity and advance our understanding of the mechanisms underlying infrared neuromodulation. This work lays a foundation for the future use of infrared light as a precise and minimally invasive tool for cortical circuit manipulation.

## 1. Introduction

Neurostimulation techniques have transformed both neuroscience research and clinical therapy by enabling precise modulation of neural activity to probe brain function and treat neurological disorders [1], [2], [3], [4]. Conventional methods, such as electrical stimulation, though effective, are constrained by factors including invasiveness, spatial specificity, and stimulation-induced electrical artefacts that interfere with neural recordings. These challenges have motivated the development of alternative approaches that are less invasive and offer superior spatial and temporal precision.

Infrared neural stimulation (INS) has emerged as a promising neuromodulation technique. It uses pulsed or continuous infrared light, typically within the 1400–2100 nm wavelength range, to evoke neuronal activity through highly controlled, localized temperature transients [5], [6]. INS utilizes the strong absorption of infrared light by water in biological tissues, leading to rapid and spatially confined heating of neural membranes and subsequent depolarization, without physical contact [7], [8]. This approach allows high spatial resolution in controlling neural activation. Furthermore, unlike electrical stimulation, INS is free from stimulation-induced electrical artifacts, enabling artifact-free electrophysiological recordings. Another significant benefit of INS is that it can be applied directly to native neural tissue without the need for prior genetic manipulation. This is in contrast, for example, to optogenetic stimulation methods, where light-sensitive opsins are expressed in the tissue [3]. This provides enhanced ease and flexibility for both experimental and clinical applications.

The biophysical mechanisms underlying INS involve spatiotemporally confined heating of neural membranes, which alters membrane properties and drives depolarization [9], [10]. Rapid changes in membrane capacitance generate a depolarizing inward current [9], [11], [12], [13]. This change may be attributed to temperature-induced modifications in membrane thickness or the formation of transient nanopores [12], [14]. Even though nanoporation has not been the focus of many INS studies, it has been shown to allow ion flux, which can depolarize neurons or activate intracellular signaling pathways [14]. Furthermore, INS-induced heating has been shown to generate stress relaxation waves, originating at the site of infrared (IR) absorption in the surrounding fluid, thereby mechanically stimulating sensory cells such as cochlear hair cells [15], [16], [17].

Temperature-sensitive ion channels, particularly those of the transient receptor potential vanilloid (TRPV) family, play a crucial molecular role in INS [18], [19]. These channels are extensively expressed throughout the peripheral and central nervous systems [20]. The discovery of heat-sensitive ion channels can be traced back to the late 1990s, when Caterina and colleagues first described TRPV1 [21]. TRPV1, also known as the capsaicin receptor, is a nonselective cation channel permeable to Na⁺, K⁺, and Ca²⁺, and is activated by various stimuli, with a critical thermal threshold above ∼43°C [22]. The role of TRPV1 in mediating responses to INS was first demonstrated by Rhee et al. [18], who showed that pulsed infrared light fails to evoke membrane depolarization in primary sensory neurons when TRPV1 channels were blocked. Similarly, Suh and colleagues [19] reported that auditory nerve action potentials could not be evoked by infrared stimulation in TRPV1 knockout mice. Other members of the TRPV family, including TRPV2, TRPV3, and TRPV4, also contribute to thermal responses but display distinct activation thresholds [23], [24], [25]. Recent research has utilized the thermo-responsiveness of the TRPV1 channel for precise and minimally invasive neuromodulation [26]. For instance, it has been demonstrated that biodegradable polymer micelles exhibit strong photothermal conversion efficiency under infrared irradiation, leading to the activation of TRPV1 channels *in vitro* [26]. This, in turn, results in a significant influx of calcium and membrane depolarization, without reaching temperatures that could be considered harmful. Similarly, TRPV1-targeting nanoplatforms that combine NIR-triggered nitric oxide release and chemodynamic therapy have shown enhanced anticancer efficacy and safety [27].

The versatility of INS has been demonstrated across diverse neuroscience applications [6], [28], [29], [30], [31]. In the central nervous system, INS can evoke intrinsic optical signals and modulate neuronal firing patterns with high spatial and temporal precision [28]. In non-human primates, it facilitates the selective activation of specific regions within the visual cortex [6]. In the auditory system, INS has been optimized for cochlear stimulation, with efficacy depending on wavelength and pulse shape [29]. It was further demonstrated that INS can facilitate both excitatory and inhibitory neuromodulation, thereby expanding its therapeutic potential [30]. In comparison with electrical stimulation, INS offers contact-free delivery, high selectivity, and compatibility with advanced imaging modalities [31], [32].

Our previous work using a neural probe with an integrated platinum heater showed that resistive heating can modulate neural activity similarly to IR irradiation [33]. To study the effects of INS on neocortical neurons, we developed an implantable photonic device delivering spatially confined IR light while simultaneously recording electrophysiological responses and monitoring temperature [34], [35]. Furthermore, we refined a Michigan-type silicon microprobe for effective INS [36] and demonstrated that low-energy continuous-wave IR light (<13 mW) can reversibly excite or suppress rat cortical neurons in a cell-type-specific manner [37]. Histological analyses confirmed the safety of repeated intracortical suppression at these low power levels [35].

Although INS has been studied extensively in *in vitro* and *ex vivo* preparations, there is still a scarcity of *in vivo* investigations, particularly in genetically modified animal models. This gap limits our understanding of the molecular mechanisms underlying the effects of INS. Genetically modified mice, such as TRPV1 knockouts, provide an opportunity to directly investigate the role of TRPV1 in INS-mediated neuromodulation. Building on our previous work [37], the present study investigates neural responses to both pulsed and continuous-wave INS in wild-type and TRPV1 knockout mice. Using high-density Neuropixels recordings, we analyzed local field potentials as well as single- and multi-unit activity, complemented by histological validation. This integrative approach allowed us to characterize the *in vivo* effects of infrared stimulation on cortical activity and to assess the contribution of TRPV1 to INS-induced responses, thereby linking channel-specific mechanisms to the observed neuromodulatory outcomes.

## 2. Materials and Methods

### 2.1. Animal surgery

All experiments were conducted in accordance with the EC Council Directive of September 22, 2010 (2010/63/EU), and all procedures were reviewed and approved by the Animal Care Committee of the HUN-REN Research Centre for Natural Sciences and by the National Food Chain Safety Office of Hungary (license number: PE/EA/01229-6/2023). Acute experiments were performed on adult male TRPV1 knockout mice (TRPV1^-/-^, purchased from Jackson Laboratory (Bar Harbor, ME, USA); bred as homozygotes; n = 5; weight: 24.80 ± 0.45 g, mean ± SD) and their genetically unaltered C57BL/6 controls (wild-type, WT; n = 5; weight: 23.80 ± 1.10 g; all male). Experiments were performed during the subjective nighttime phase of the animals.

The surgery protocol followed that described in Horváth et al. [38]. Briefly, anesthesia was induced by intraperitoneal injection of a ketamine (100 mg/kg body weight) and xylazine (10 mg/kg body weight) mixture. To maintain the surgical plane of anesthesia during experiments, supplementary intramuscular injections of ketamine/xylazine were administered at regular intervals (1 – 2 injections per hour). Body temperature was kept at physiological levels with a homeothermic heating pad controlled by a temperature controller unit (Supertech, Pécs, Hungary).

Once surgical anesthesia was achieved, animals were placed in a stereotaxic frame (David Kopf Instruments, Tujunga, CA, USA), and the skin and connective tissue were removed to expose their skull. Next, using a dental drill, a circular craniotomy (∼2 mm in diameter) was prepared over the target region in the left hemisphere. First, an optical fiber with a diameter of 125 µm was inserted into the trunk region of the somatosensory cortex (S1Tr) at a 30° angle relative to vertical, followed by implantation of a single-shank Neuropixels 1.0 silicon probe within <300 µm of the fiber [39]. Target coordinates of the optical fiber were anteroposterior (AP): -1.6 mm and mediolateral (ML): 1.8 mm, while the Neuropixels probe was implanted at approximately AP: -1.85 mm and ML: 1.55 mm relative to the bregma [40]. The dura mater was left intact during the implantation procedure.

The optical fiber and the silicon probe were mounted on separate motorized stereotaxic micromanipulators (Robot Stereotaxic, Neurostar, Tübingen, Germany) and lowered into the brain at a slow speed (∼2 μm/s; [41]) to depths of 0.6 mm (layer 5 of S1Tr) and 3.5 mm, respectively. To prevent tissue dehydration, room-temperature saline was regularly applied to the craniotomy. A stainless-steel needle inserted into the nuchal muscle of animals served as both an external reference electrode and the ground.

### 2.2. Optical stimulation

Infrared light was delivered through the optical fiber, generated by a fiber-coupled laser diode (LPSC-1550-FG105LCA-SMA, Thorlabs GmbH, Germany) at 1550 nm, a wavelength chosen based on previous studies [34], [35], [37]. The laser had a nominal maximum output power of 80 mW at the exit side of the optical fiber.

The IR stimulation protocol consisted of alternating 4-minute ON (laser active) and 4-minute OFF (laser inactive) periods, allowing the stimulated region to return to baseline temperature between stimulation cycles, as validated in our previous work [37]. Each recording started with a 2-minute period containing spontaneous cortical activity. Earlier histological work confirmed that repeated application of these stimulation parameters does not alter cortical cell density, supporting the safety and viability of this approach [34], [35].

Both continuous-wave (CW) and pulsed (500 Hz; rectangular pulses, 50% duty cycle) illumination modes were applied in all mice, with the order of presentation randomized across animals. Five consecutive trials (8-min ON-OFF cycles) were performed for each stimulation mode. The IR laser diode was operated at 378 mA for pulsed stimulation and at 200 mA for CW stimulation, both yielding an ∼5°C temperature increase. Current delivery was controlled by a Keithley 2611B SYSTEM SourceMeter (Keithley Instruments Inc., OH, USA) and the same device was used to generate synchronized trigger signals to ensure precise stimulation timing.

Thermal effects of IR stimulation were assessed *in vitro* using a calibrated platinum temperature sensor, following established methods [35], [37]. Temperature changes measured relative to baseline varied slightly with IR stimulation mode: 4.9 °C for pulsed (500 Hz) and 5.1 °C for CW stimulation.

### 2.3. Electrophysiological recorings

As in our previous study [37], cortical electrical activity, including both local field potentials (LFPs) and spiking activity, was recorded using the Neuropixels recording system connected to a PXIe-1071 chassis equipped with an MXI-Express interface (National Instruments, Austin, Texas, USA). Neural signals were acquired from 384 channels selected out of 960 available recording sites. Spiking activity was sampled at 30 kHz per channel, and LFPs at 2.5 kHz per channel, both with 10-bit resolution. Data acquisition was performed using the SpikeGLX software package (https://billkarsh.github.io/SpikeGLX/).

For data collection, only the first 384 recording sites located closest to the probe tip (bank0), were used. For each mouse, two recordings were obtained, one for each stimulation mode (CW and 500 Hz pulsed). The duration of each recording was 42 minutes. Electrophysiological data were obtained from the somatosensory cortex, hippocampus, and thalamus; however, the present study focused exclusively on the analysis of somatosensory cortical activity, as the optical fiber was implanted in the cortex and the heating effect of the IR light is spatially restricted to approximately 1 mm.

### 2.4. RNAscope in situ hybridization

Due to the lack of a reliable antibody, we used the highly selective RNAscope in situ hybridization (ISH) technique for the detection of the TRPV1 mRNA. Experiments were performed on 4-month-old male WT mice (n = 4). Animals were deeply anesthetized with an overdose of urethan (2.4 g/kg) and perfused transcardially with ice-cold 0.1 M phosphate-buffered saline (PBS, pH = 7.4), followed by 4% paraformaldehyde (PFA) in Millonig’s phosphate buffer. Brains were then dissected and post-fixed in PFA solution at 4 °C for 72 h. Coronal brain sections (30 mm thickness) were cut using a vibrating microtome (VT1000S, Leica Biosystems, Wetzlar, Germany) and stored in 1x PBS containing 0.01% Na-azide (Merck KGaA, Darmstadt, Germany) until use.

RNAscope ISH was carried out according to the manufacturer’s protocol using RNAscope Multiplex Fluorescent Reagent Kit v2 (Advanced Cell Diagnostic (ACD), Newark, CA, USA). Brain sections containing the somatosensory cortex were hybridized with probes specific for mouse *Trpv1* (ACD, Cat. No. 313331), mouse *Vglut1* (ACD, Cat. No. 416631) and mouse *Gad1* (ACD, Cat. No. 400951). The probes were visualized with fluorescein (1:750 for *Vglut1*), cyanin 3 (Cy3; 1:750 for *Trpv1*) and cyanin 5 (Cy5; 1:750 for *Gad1*).

A mouse 3-plex positive control probe (ACD, Cat. No. 320881), targeting *Polr2a* mRNA (fluorescein), *Ppib* mRNA (Cy3), and *Ubc* mRNA (Cy5) showed a clear signal in randomly selected regions of interest. Conversely, no signal was detected with the 3-plex negative control probe (ACD, Cat.No. 320871) after hybridization and channel development (images not shown).

All sections were counterstained with 4’,6-diamidino-2-phenylindole (DAPI) and mounted with ProLong Glass Antifade Mountant (Thermo Fisher Scientific, Waltham, MA, USA) for confocal imaging. Preparations were scanned using a Nikon Eclipse Ti2-E confocal microscope with a 60× objective. Fluorescent signals were pseudocolored as follows (see Fig. 4); DAPI (blue), *Vglut1* (green), *Trpv1* (red) and *Gad1* (white). Brightness and contrast adjustments, including maximum intensity projections of individual channels, were performed using FIJI (v1.53c, NIH, USA; [42]).

### 2.5. Data analysis

#### 2.5.1. Spike sorting and single unit curation

Spikes from Neuropixels recordings were detected and sorted using KiloSort (v3.0) [43], with the spike sorting algorithm executed on an NVIDIA V100 GPU. Spike sorting was applied to the two recordings (pulsed and CW stimulation) separately. Following automated sorting, single units were manually curated based on their autocorrelograms, crosscorrelograms, firing rates, amplitude distributions, and spike waveform features.

Additional units were excluded from analysis based on quality metrics computed with SpikeInterface (v0.98.2). The quality metrics employed were the presence ratio, interspike interval (ISI) violations, and amplitude cutoff. The presence ratio, ranging from 0 to 1, quantifies the proportion of the recording during which a unit is active, with higher ratios indicating stable and reliable firing throughout the session. For the presence ratio, a threshold of 0.9 was applied, retaining all units above this value. ISI violations measure the rate of refractory period violations, with lower values reflecting minimal contamination from other clusters; units exceeding a threshold of 2 were excluded. The amplitude cutoff, ranging from 0 to 0.5, assesses the miss rate based on amplitude histograms. Units with amplitude cutoff values above 0.01 were removed from further analysis.

#### 2.5.2. Differentiation between cortical cell types

Within the neocortex, neurons can be broadly classified into two main types: inhibitory interneurons and excitatory principal cells [44]. The distinction is based on characteristic features of their extracellular action potential waveforms and firing patterns, including measures related to burst firing such as ACG_tau_rise [45]. This measure reflects the proportion of spikes occurring at short interspike intervals, providing an estimate of a neuron’s burst tendency based on its autocorrelogram. Interneurons can be further divided into wide-spiking and narrow-spiking subtypes according to their waveform properties. Principal cells, which display wide action potentials, typically have shorter ACG_tau_rise values consistent with their burst-prone firing, whereas wide-spiking interneurons usually show longer ACG_tau_rise values [45].

In our recordings, Hartigan’s dip test confirmed that the distribution of spike durations (trough-to-peak times) was bimodal (p = 0.031), indicating a clear distinction between the wide-spiking and narrow-spiking populations. To identify the three putative neuron types (pyramidal cells, narrow-spiking interneurons, and wide-spiking interneurons), CellExplorer (v1.2) was used [45]. This software automatically classifies the extracted single units into one of the three groups based on spike waveform features (trough-to-peak times) and autocorrelogram characteristics (rise component of the triple exponential function fit to the autocorrelogram; ACG_tau_rise [45]). Following automated classification, each unit was manually inspected in CellExplorer, using various features such as firing rate to correct any misclassifications.

#### 2.5.3. Identification of the laminar location of single units

The laminar position of recording sites was determined using LFP- and spiking activity-related features. Due to the thinness of layers 2 and 4 in mice and the difficulty of precisely delineating their boundaries, single units from superficial and input layers (layers 1-4) were grouped. Under ketamine/xylazine anesthesia, superficial layers exhibit sparse spontaneous population activity and fewer active neurons compared to infragranular layers (layers 5 and 6; [46], [47]). In contrast, layer 5 shows the strongest neuronal activity and contains the cell bodies of the largest pyramidal cells in any cortical layer. Layer 5 was identified based on multi-unit activity (MUA) amplitudes during up-states and its depth relative to the cortical surface [48]. Layer 6 was defined as the region below layer 5, where MUA amplitude decreases toward the bottom of the cortex and short-duration positive spikes appear near the transition to the underlying white matter [48]. Each recording site was assigned to one of these cortical layers, and individual cortical single units were mapped to the recording site where their mean spike waveform exhibited the largest amplitude.

To ensure cross-animal comparability for visualization purposes, depth normalization was performed to account for variations in recording channel labeling between animals, following established literature and our previous methodological paradigm [37], [49], [50].

#### 2.5.4. Firing rate analysis

For each single unit, the relative firing rate change compared to firing rates measured during the baseline period was calculated in ten-second intervals during each IR stimulation (ON) period, as described in Equation *(1)*:

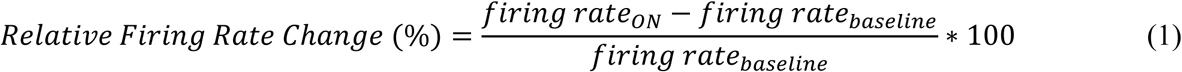

Based on our definition, for example, 0% indicates no change in the firing rate of the particular unit relative to the baseline, while 100% corresponds to a doubling of the firing rate. For each stimulation trial, the baseline firing rate was defined as the average firing rate during the last two minutes of the recovery (OFF) period preceding stimulation (or, for the first stimulation trial, the first two minutes of the recording), representing spontaneous cortical activity between stimulation periods. Instead of using a single baseline period for all stimulation trials (e.g., the first two minutes of the recording containing spontaneous activity), we employed multiple, local baseline periods to minimize the influence of global brain state fluctuations unrelated to stimulation. Although baseline firing rates gradually increased across consecutive stimulation trials - reaching a plateau around the fourth trial (Supplementary Fig. S1) - suggesting that repeated stimulation may induce prolonged changes in baseline neuronal firing, this effect was not further investigated in this study. This modulation was more pronounced during 500 Hz stimulation, consistent with our previous findings in anesthetized rats [37].

To remove outliers resulting from inaccurate spike detections, single units with relative firing rate changes exceeding 500% were first excluded. After this, we calculated thresholds according to the distribution of baseline firing rates: the lower threshold was defined as (10^th^ percentile - 3x interpercentile range), and the upper threshold as (90^th^ percentile + 3x interpercentile range), where the interpercentile range is the difference between the 90^th^ and 10^th^ percentiles. Single units with baseline firing rates below the lower threshold or above the upper threshold were excluded. These two filtering steps resulted in the exclusion of approximately 3% of units from WT animals (n = 26 units) and ∼7% of units from KO animals (n = 61 units).

Next, single units were classified based on relative firing rate changes into three categories: increased (firing rate increased to IR stimulation), suppressed (firing rate decreased to stimulation), or unaffected (no significant change in firing rate). For each neuron, the firing rate change during stimulation (24 firing rate change values computed from ten-second intervals of the four-minute stimulation trial) was tested against zero (baseline firing rate) using Wilcoxon signed-rank test [51], [52]. Neurons were considered significantly modulated (activity increased or suppressed) if p < 0.1. Single units were classified as increased or suppressed if they showed a significant increase or decrease in firing rate, respectively, in at least 3 out of 5 trials (60%). Neurons with non-significant firing rate changes (p ≥ 0.1 for more than 2 trials) were considered unaffected. Firing rate changes of all units were also visually inspected and manually curated to ensure accurate classification.

To assess the temporal dynamics of neuronal responses to IR stimulation, the following two latency parameters were measured for each suppressed or increased single unit: rise time, defined as the time at which activity reached 90% of the maximal relative firing rate change during the ON periods, and fall time, defined as the time at which activity decreased to 10% of this firing rate change during the subsequent OFF periods.

#### 2.5.5. Local field potential analysis

To analyze LFPs, raw LFP signals from cortical channels were processed. To reduce line noise, notch filters were applied at 50 Hz and its harmonics (100 Hz and 150 Hz). After preprocessing, the LFP was bandpass filtered (4th-order Butterworth, zero-phase shift) within well-established frequency ranges corresponding to various brain rhythms: delta (0.5–4 Hz), theta (4–8 Hz), alpha (8–12 Hz), beta (12–30 Hz), and gamma (30–100 Hz) bands.

We estimated the power in each frequency band using the *bandpower* function of MATLAB. For every recording, the analysis focused on the last minute of all stimulation (ON) and recovery periods (used as baseline). Within each 60-second window, the average spectral power was derived across channels and trials for each frequency band. To allow comparison between wild-type and knockout mice, we computed the change in LFP power relative to the baseline period using Equation (2):

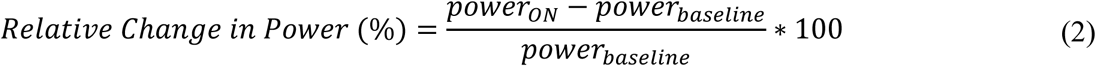

#### 2.5.6. State detection and multi-unit activity analysis

To extract MUA, raw signals were bandpass filtered between 500 and 5000 Hz using a third-order Butterworth filter (zero-phase shift). The filtered signals were then rectified and downsampled by a factor of 10, resulting in an effective sampling rate of 3 kHz. Next, MUA signals from all cortical channels were summed to represent cortical population activity on a single channel and to capture slow network state fluctuations. This summed signal was then lowpass filtered at 30 Hz (third-order, zero-phase shift Butterworth filter), resulting in the smoothed population activity (SPA) signal. To detect the onset of up- and down-states, a threshold was computed based on the SPA amplitude within putative down-states (*mean* + 3 ∗ *standard deviation* [39], [54]). Up- and down-state onsets were identified by threshold crossings, with enforced minimum durations (for up-states: 100 ms; for down-states: 50 ms) to reduce false detections. The duration of each detected state was calculated to characterize state length distributions. Up- and down-states with excessively short or long durations (likely resulting from inaccurate state-onset detections) were excluded (±3σ).

Next, we calculated averages from the SPA signal (reflecting MUA) aligned to the start of up-states for both stimulation (ON) and baseline (BL) periods. Short segments were extracted from the SPA signal around each detected up-state onset (from 150 ms before to 500 ms after the onset). For each stimulation trial (separately for ON and BL periods), these segments were then averaged across up-states. Resulting segments were subsequently averaged across trials. Only up-states detected during the last minute of the ON (stimulation) and OFF (baseline) periods were used to compute SPA averages. Finally, we calculated the average SPA amplitude (referred to as MUA amplitude in the Results section) within the 10–100 ms window following the up-state onset, both during stimulation and the baseline period. The change in MUA amplitude induced by IR stimulation, relative to baseline, was quantified using Equation (3):

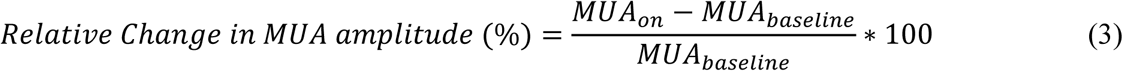

### 2.6. Statistical analysis

All statistical analyses were performed using GraphPad Prism (v9.5.1). Normality of the data was assessed using D’Agostino & Pearson and Shapiro-Wilk tests. For comparisons between two groups, paired or unpaired Student’s t-tests were applied to normally distributed data, while the Wilcoxon signed-rank test or Mann–Whitney U test was used for non-normal data. For comparisons involving more than two groups, nonparametric Friedman or Kruskal–Wallis tests were employed, as the data did not follow a normal distribution. When these tests indicated a significant effect, Dunn’s post hoc test was applied to evaluate pairwise group differences. Statistical significance was defined as p < 0.05. Values in the Results section are reported as mean ± standard deviation, unless stated otherwise.

## 3 Results

### 3.1 Effects of infrared stimulation on spontaneous cortical activity in anesthetized wild-type mice

We first investigated how infrared stimulation influences spontaneous neural activity in the primary somatosensory cortex of wild-type mice (n = 5). Cortical activity was recorded *in vivo* under ketamine/xylazine anesthesia using a single-shank Neuropixels silicon probe positioned adjacent to an optical fiber used for stimulation (Fig. 1A). Pulsed (500 Hz) or continuous-wave (CW) IR light, with a wavelength of 1550 nm, was delivered through the tip of the fiber placed in infragranular layers. The stimulation paradigm consisted of alternating four-minute periods of IR stimulation and recovery (five repetitions). During IR stimulation, cortical temperature increases by several degrees Celsius, whereas recovery periods allow the tissue to return to baseline temperature (Fig. 1B).

**Figure 1.**
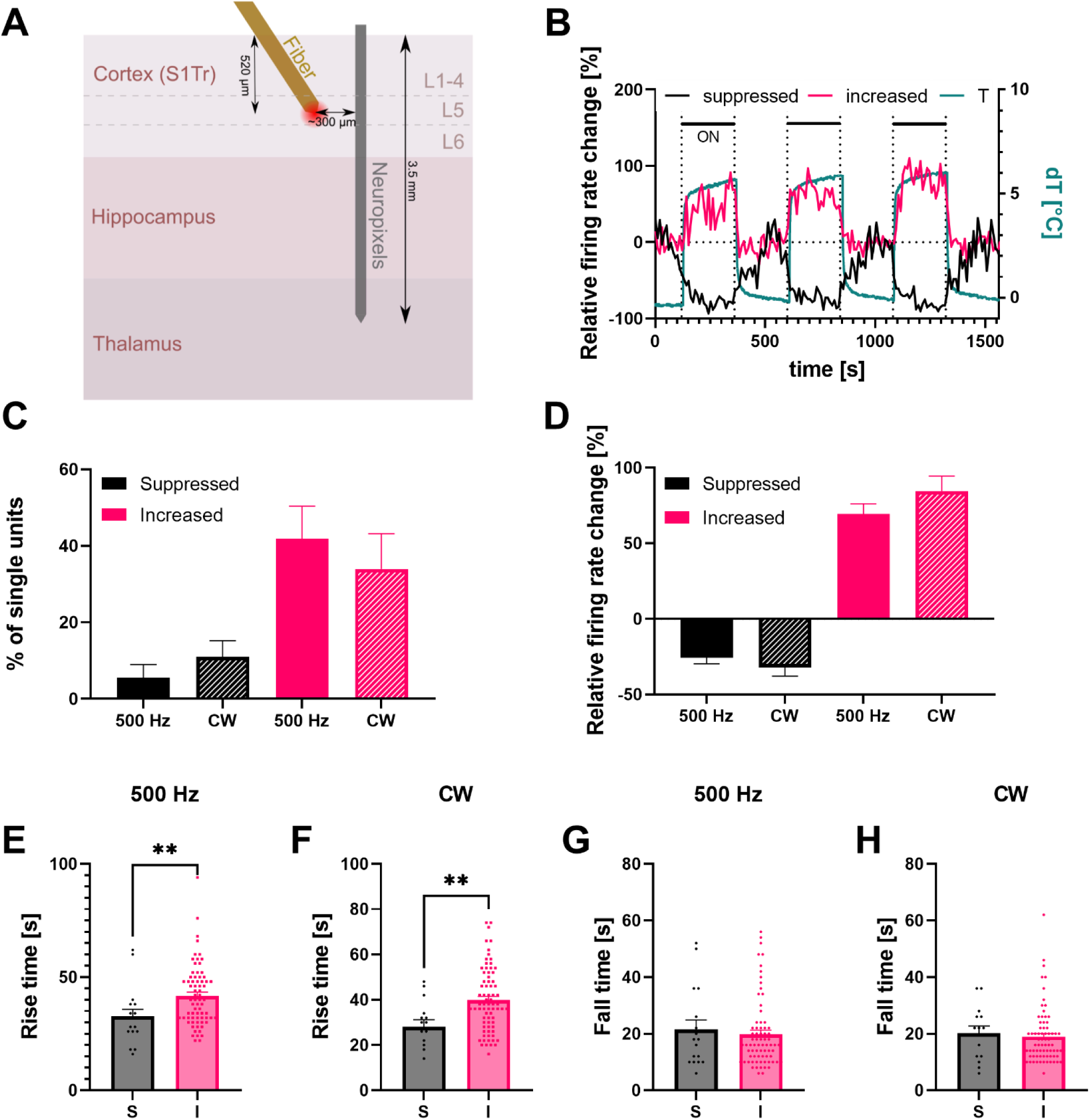
Experimental setup and effects of infrared stimulation on single-unit activity in anesthetized wild-type mice. (A) Schematic of the experimental setup (not to scale). S1Tr, trunk region of the primary somatosensory cortex; L1-4, Layers 1-4; L5, Layer 5; L6, Layer 6. (B) Example traces of two single units (firing rate change relative to baseline) exhibiting increased (red) or suppressed (black) activity during continuous-wave (CW) stimulation. The corresponding temperature change measured in vitro during CW stimulation is shown in green. Black horizontal bars indicate stimulation periods (ON). Three out of five trials are displayed. (C) Proportion of single units exhibiting suppressed (black) or increased (red) activity during pulsed (500 Hz) and CW stimulation. (D) Average relative firing rate changes in neurons with increased (red) or suppressed (black) activity during pulsed (500 Hz) and CW stimulation. (E–F) Rise times of single units with suppressed (S) and increased (I) activity during pulsed (E) and CW (F) stimulation. Rise times were significantly longer in neurons with increased activity. ** p < 0.01. (G–H) Fall times of single units with suppressed (S) and increased (I) activity during 500 Hz (G) and CW (H) stimulation. Error bars in all panels represent the standard error of the mean.

To assess the effects of IR stimulation, we first analyzed changes in the properties of single-unit activity, including the proportion of neurons whose firing was modulated by IR light, alterations in firing rates relative to recovery periods, and the dynamics of firing rate changes that reflect how rapidly IR stimulation modulates activity. We further compared these properties across major cortical cell types, that is, inhibitory interneurons and excitatory principal cells. In addition, we investigated whether the distribution of units responsive to IR stimulation varied across cortical layers. Finally, given that slow-wave activity is the dominant neural rhythm under ketamine/xylazine anesthesia in the cortex [48], and that IR stimulation has been shown to modulate this oscillation in rats [54], we investigated whether similar effects occur in the somatosensory cortex of mice. For all analyses, we compared the effects of the two stimulation modes, namely 500 Hz and CW stimulation.

#### 3.1.1 Infrared stimulation-related changes in single unit properties

Spike sorting was performed on the raw recordings to isolate single-unit activity, followed by the application of various quality metrics to exclude low-quality units. Across all animals, a total of 370 single units were extracted (74.0 ± 56.6 units per mouse). Comparable numbers of units were obtained for both stimulation modes (500 Hz: 37.0 ± 27.7 units per recording; CW: 37.0 ± 31.5 units). Next, for each neuron, we calculated the relative firing rate change during IR stimulation with respect to the baseline period preceding stimulation.

In response to infrared stimulation, firing of single units was either suppressed (Fig. 1B, black), increased (Fig. 1B, red), or unaffected. Changes in firing rates were correlated with temperature fluctuations measured during stimulation and recovery periods *in vitro* (Fig. 1B, green). Approximately 8% of neurons (n = 31/370 units) displayed suppressed activity, while around 40% (n = 147/370 units) showed an increase in firing rate in response to stimulation (Fig. 1C). The majority of neurons remained unaffected by IR light (∼52%, n = 192/370 units; Fig. 1C). Differences in proportions between pulsed and CW stimulation modes were not significant, although CW stimulation elicited a slightly higher fraction of units with suppressed activity and a slightly lower fraction of units with increased firing rates compared to 500 Hz stimulation (CW vs. 500 Hz; suppressed: 10.92% ± 9.49% vs. 5.56% ± 7.56%, Wilcoxon signed-rank test, p = 0.625; increased: 33.95% ± 20.60% vs. 41.87% ± 19.03%, Wilcoxon signed-rank test, p = 0.8125; Fig. 1C).

Infrared stimulation decreased the firing rate of suppressed units by approximately 30% and elevated the firing rates of units with increased activity by about 80% (Fig. 1D). Suppressed units exhibited a slightly greater decrease in firing rates during CW stimulation compared to 500 Hz stimulation (CW vs. 500 Hz; - 32.02% ± 21.85% vs. -25.73% ± 16.13%, Student’s t-test, p = 0.3640; Fig. 1D), and units with increased activity showed a slightly larger increase; however, these differences were not statistically significant (CW vs. 500 Hz; 84.57% ± 84.98% vs. 69.59% ± 54.69%, Mann-Whitney U test, p = 0.8309; Fig. 1D). These trends may indicate a potentially stronger effect of continuous-wave stimulation relative to pulsed stimulation.

Next, we examined the temporal dynamics of single unit firing in response to IR stimulation. For each unit, we computed the time required to reach 90% of the maximum firing rate change during the ON period (rise time) for both pulsed (Fig. 1E) and CW stimulation (Fig. 1F). For both stimulation modes, rise times were significantly longer in neurons with increased activity compared to those with suppressed activity (suppressed vs. increased; 500 Hz: 32.70 s ± 12.76 s vs. 41.75 s ± 13.49 s, Mann–Whitney U test, p = 0.0094; CW: 28.00 s ± 12.49 s vs. 39.86 s ± 14.20 s, Mann–Whitney U test, p = 0.005; Fig. 1E and F). Rise times did not differ significantly between the two stimulation conditions, although firing rate changes were slightly faster during CW stimulation (CW vs. 500 Hz; suppressed: 28.00 s ± 12.49 s vs. 32.7 s ± 12.76 s, Mann-Whitney U test, p = 0.4481; increased: 39.86 s ± 14.20 s vs. 41.75 s ± 13.49 s, Mann-Whitney U test, p = 0.4289).

We also computed the time required to reach 10% of the maximum firing rate change after the end of IR stimulation (fall time) for both stimulation modes (Fig. 1G and H). We found no significant differences in fall times between suppressed and increased neurons (suppressed vs. increased; 500 Hz: 21.41 s ± 14.12 s vs. 19.86 s ± 12.13 s, Mann–Whitney U test, p = 0.8075; CW: 16.15 s ± 8.85 s vs. 18.97 s ± 9.89 s, Mann–Whitney U test, p = 0.5568; Fig. 1G and H) or between pulsed and CW stimulation (CW vs. 500 Hz; suppressed: 20.14 s ± 9.78 s vs. 21.41 s ± 14.12 s, Mann–Whitney U test, p = 0.9471; increased: 18.43 s ± 8.43 s vs. 19.86 s ± 12.13 s, Mann–Whitney U test, p = 0.6840). Rise and fall times were consistent across trials, indicating stable temporal dynamics throughout repeated stimulation cycles (Supplementary Fig. S2).

Based on electrophysiological features, single units can be separated into interneurons and principal cells [44], [45]. To examine neuron type-specific effects of IR stimulation, we classified the isolated units as putative narrow-spiking interneurons, wide-spiking interneurons, or principal cells, using spike duration and burstiness measures (Fig. 2A and B; see Methods for details). In wild-type mice, approximately one-third of units were narrow-spiking interneurons (N-I), ∼9% were wide-spiking interneurons (W-I), and the remaining slightly more than half units were classified as principal cells (P; Fig. 2C).

**Figure 2.**
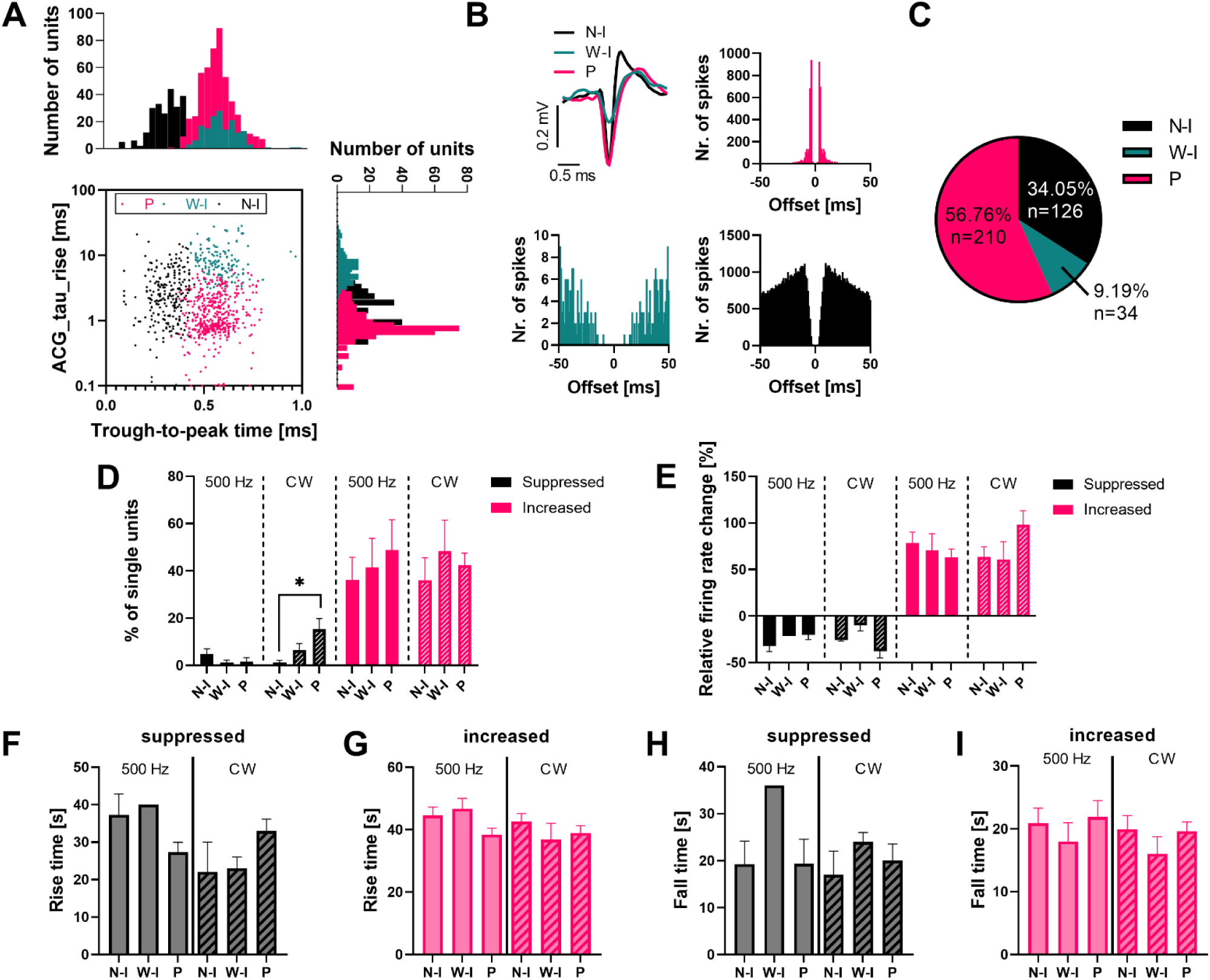
2 Neuron type-dependent effects of infrared stimulation in the somatosensory cortex of wild-type mice. (A) Electrophysiological features used to separate the three neuron types and neuron-type specific distribution of single units. P, principal cells; N-I, narrow-spiking interneurons; W-I, wide-spiking interneurons. (B) Representative mean spike waveforms of a narrow-spiking interneuron (black), a wide-spiking interneuron (green), and a principal cell (red), with corresponding autocorrelograms. (C) Proportions of the three neuron types (n = 370 single units in total). Percentages and cell counts are indicated for each type. (D) Proportion of single units showing suppressed (black) or increased (red) activity during pulsed (500 Hz) and continuous-wave (CW) stimulation, grouped by neuron type. The ratio of suppressed principal cells was significantly higher than that of narrow-spiking interneurons. * p < 0.05. (E) Relative firing rate change of units with suppressed and increased activity, broken down by neuron type. (F–G) Rise times of the three cell types with suppressed (F) or increased (G) activity during 500 Hz and CW stimulation. (H–I) Fall times of the three neuron types with suppressed (H) and increased (I) activity during 500 Hz and CW stimulation. Error bars in all panels indicate the standard error of the mean.

The proportion of units with increased activity was similar across cell types for both stimulation modes (N-I vs. W-I vs. P; 500 Hz: 36.22% ± 21.19% vs. 41.32% ± 27.71% vs. 48.82% ± 28.73%, Friedman test, p = 0.2284; CW: 36.03% ± 21.24% vs. 48.24% ± 29.65% vs. 42.46% ± 11.09%, Friedman test, p = 0.0849; Fig. 2D; Supplementary Table S1). The proportion of suppressed neurons was also similar, although the fraction of putative principal cells was significantly higher than that of narrow-spiking interneurons during CW stimulation (N-I vs. W-I vs. P; 500 Hz: 4.69% ± 5.24% vs. 1.11% ± 2.48% vs. 1.63 % ± 3.65%, Friedman test, p = 0.1111; CW, 1.32% ± 1.81% vs. 6.48% ± 6.13% vs. 15.29% ± 10.13%, Friedman test, p = 0.0216; Fig. 2D).

When examining relative firing rate changes in suppressed units, no statistically significant differences were observed across cell types for either stimulation mode (N-I vs. W-I vs. P; 500 Hz: -32.09% ± 16.78% vs. - 21.59% ± 0.00% vs. -19.88% ± 15.10%, Kruskal-Wallis test, p = 0.2109; CW: -25.82% ± 1.20% vs. -9.94% ± 8.58% vs. -37.67% ± 14.02%, Kruskal-Wallis test, p = 0.2414; Fig. 2E). Differences between units with increased activity were also not significant across cell types (N-I vs. W-I vs. P; 500 Hz: 78.79% ± 57.35% vs. 70.53% ± 50.73% vs. 63.18% ± 54.20%, Kruskal-Wallis test, p = 0.4632; CW: 63.66% ± 49.16% vs. 61.09% ± 49.82% vs. 98.28% ± 99.80%, Kruskal-Wallis test, p = 0.5431; Fig. 2E). However, CW stimulation appeared to exert a slightly stronger effect on principal cells, affecting both suppressed and increased units.

We next compared rise times across neuron types during different stimulation conditions. Overall, no significant differences were observed for both suppressed units (N-I vs. W-I vs. P; 500 Hz: 37.25 s ± 15.82 s vs. 40.00 s ± 0.00 s vs. 27.25 s ± 7.55 s, Kruskal–Wallis test, p = 0.1640; CW: 22.00 s ± 11.31 s vs. 23.00 s ± 4.24 s vs. 33.00 s ± 9.90 s, Kruskal–Wallis test, p = 0.1998; Fig. 2F) and neurons with increased firing rates (N-I vs. W-I vs. P; 500 Hz: 44.64 s ± 12.91 s vs. 46.67 s ± 10.00 s vs. 38.37 s ± 13.36 s, Kruskal–Wallis test, p = 0.0162; CW: 42.64 s ± 11.88 s vs. 36.86 s ± 13.70 s vs. 38.98 s ± 15.34 s, Kruskal–Wallis test, p = 0.3708; Fig. 2G). Although the Kruskal-Wallis test indicated a significant difference between the rise times of different neuron types with increased activity for 500 Hz stimulation, no significant differences were found in post-hoc pairwise comparisons.

Fall times were also similar across cell types and stimulation conditions for suppressed units (N-I vs. W-I vs. P; 500 Hz: 19.25 s ± 13.97 s vs. 36.00 s ± 0.00 s vs. 19.33 s ± 15.77 s, Kruskal–Wallis test, p = 0.5576; CW: 17.00 s ± 7.07 s vs. 24.00 s ± 2.83 s vs. 22.73 s ± 13.97 s, Kruskal–Wallis test, p = 0.7902; Fig. 2H) and units with increased activity (N-I vs. W-I vs. P; 500 Hz: 20.92 s ± 12.16 s vs. 18.00 s ± 8.95 s vs. 21.91 s ± 16.09 s, Kruskal–Wallis test, p = 0.8896; CW: 19.93 s ± 10.23 s vs. 16.00 s ± 7.30 s vs. 19.62 s ± 9.92 s, Kruskal–Wallis test, p = 0.5512; Fig. 2I).

Next, we assigned the isolated single units to cortical layers using distinct electrophysiological markers (see Methods for details) to study the laminar distribution of neurons affected by stimulation. Units located in layers 1-4 were grouped and are referred to as the superficial/input layers (L1/2/3/4; 15.2 ± 17.7 units per mouse, n = 76 units in total), while layers 5 (L5; 22.1 ± 23.1 units, n = 113 units in total) and 6 (L6; 36.2 ± 25.1 units, n = 181 units in total) were analyzed separately. The number of single units recorded in each cortical layer for every mouse is provided in Supplementary Table S2. We found no statistically significant differences in the proportion of single units exhibiting increased or suppressed activity across cortical layers under either stimulation mode (Fig. 3A; Supplementary Table S3). Similarly, no significant differences were found for relative firing rate changes across layers (Fig. 3B; Supplementary Table S4).

**Figure 3.**
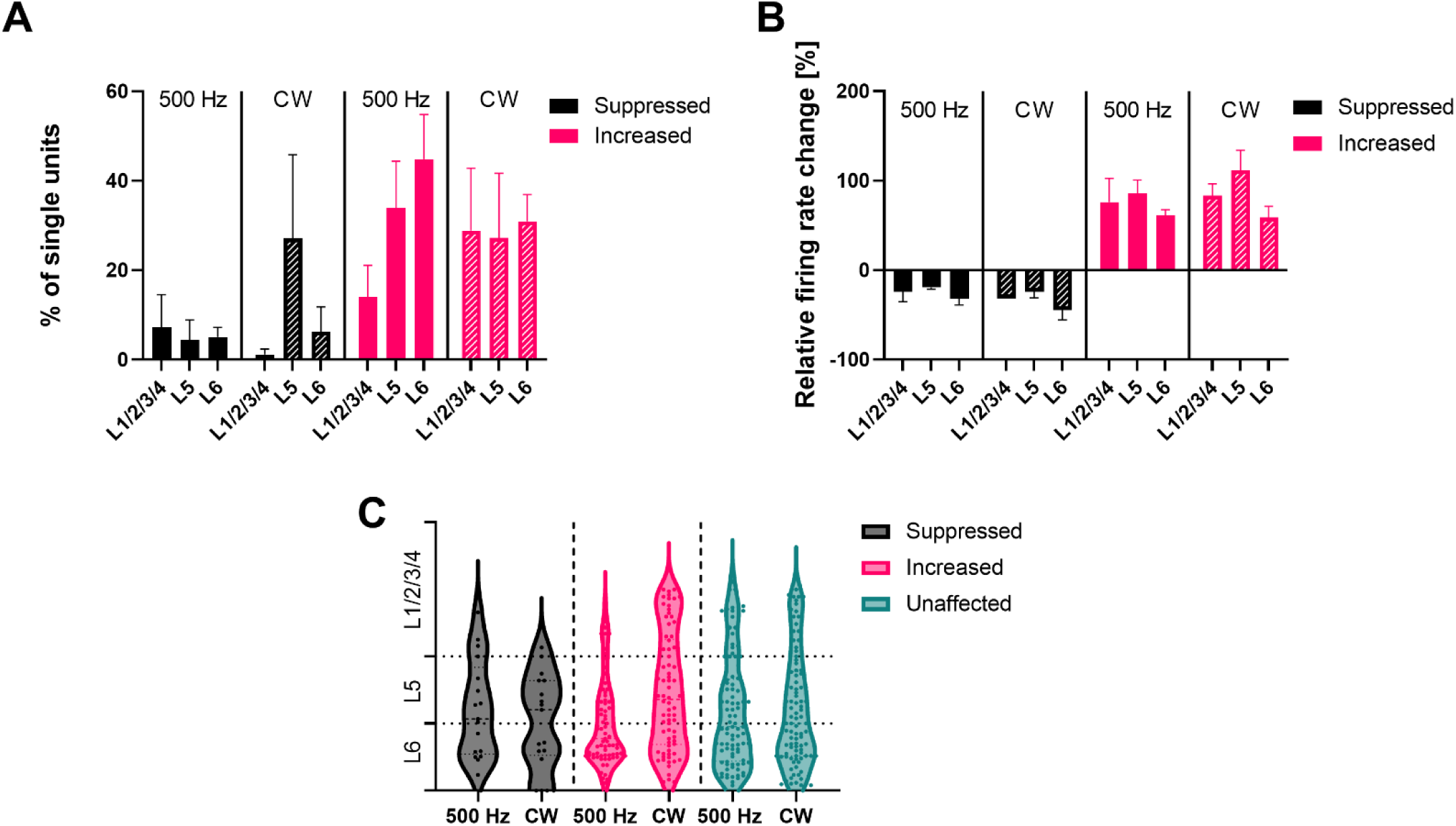
3 Laminar analysis of the effects of infrared stimulation on cortical activity in wild-type mice. (A) Proportions of single units showing suppressed (black) or increased (red) activity in the superficial and input layers (L1/2/3/4), layer 5 (L5), and layer 6 (L5) to infrared pulsed (500 Hz) or continuous-wave (CW) stimulation. (B) Average of relative firing rate change of single units with suppressed (black) or increased (red) activity in the superficial and input layers, layer 5, and layer 6 to stimulation. (C) Distribution of single units with suppressed (black), increased (red), or unaffected (green) responses to stimulation, plotted by cortical depth. Each point represents a single unit. The y-axis indicates the normalized depth of the neurons, with dashed horizontal lines marking laminar boundaries. Error bars in all panels represent the standard error of the mean.

Based on the laminar distribution of units grouped by their response to IR simulation (Fig. 3C), most units were located in the infragranular layers, consistent with earlier observations that supragranular layers exhibit sparse firing and therefore yield fewer sortable units [46], [47], [48]. Although a slightly higher number of units appeared to be affected by stimulation in layer 6 (Fig. 3A), with increased units showing smaller firing rate changes and suppressed units showing larger changes compared to the other two layer groups (Fig. 3B), these differences did not reach statistical significance. These results suggest that no single cortical layer appears to play a dominant role in IR stimulation–induced changes.

#### 3.1.2 Effects of infrared stimulation on ketamine/xylazine-induced slow-wave activity

To gain a more profound understanding of the effects of infrared stimulation on ongoing cortical dynamics, we examined potential changes in the properties of cortical slow-wave activity (corresponding to the 0.5 - 4 Hz frequency band of local field potentials) induced by ketamine/xylazine anesthesia [48]. Thus, we examined changes in spectral power of the LFP in the 0.5 - 4 Hz band (termed here delta) during stimulation and recovery periods. We found that delta power increased notably during IR stimulation compared to the baseline periods for both stimulation modes, and the difference was significant for CW stimulation (BL vs. ON; 500 Hz: 1152.96 µV^2^ ± 582.09 µV^2^ vs.1328.38 µV^2^ ± 575.03 µV^2^, paired t-test, p = 0.0737; CW: 1046.84 µV^2^ ± 610.01 µV^2^ vs. 1294.57 µV^2^ ± 769.59 µV^2^, paired t-test, p = 0.0323; Fig. 4A and B). This increase was in agreement with previous findings [53], [54]. Since slow waves group faster brain oscillations [55], we also examined changes in spectral power in higher frequency bands of the LFP, including theta (4-8 Hz), alpha (8-12 Hz), beta (12-30 Hz), and gamma (30-100 Hz) bands. The power in the theta band did not change during stimulation, while power in the alpha, beta, and gamma bands showed a slight decrease, although these changes were not significant (Supplementary Fig. S3; Supplementary Table S5).

**Figure 4.**
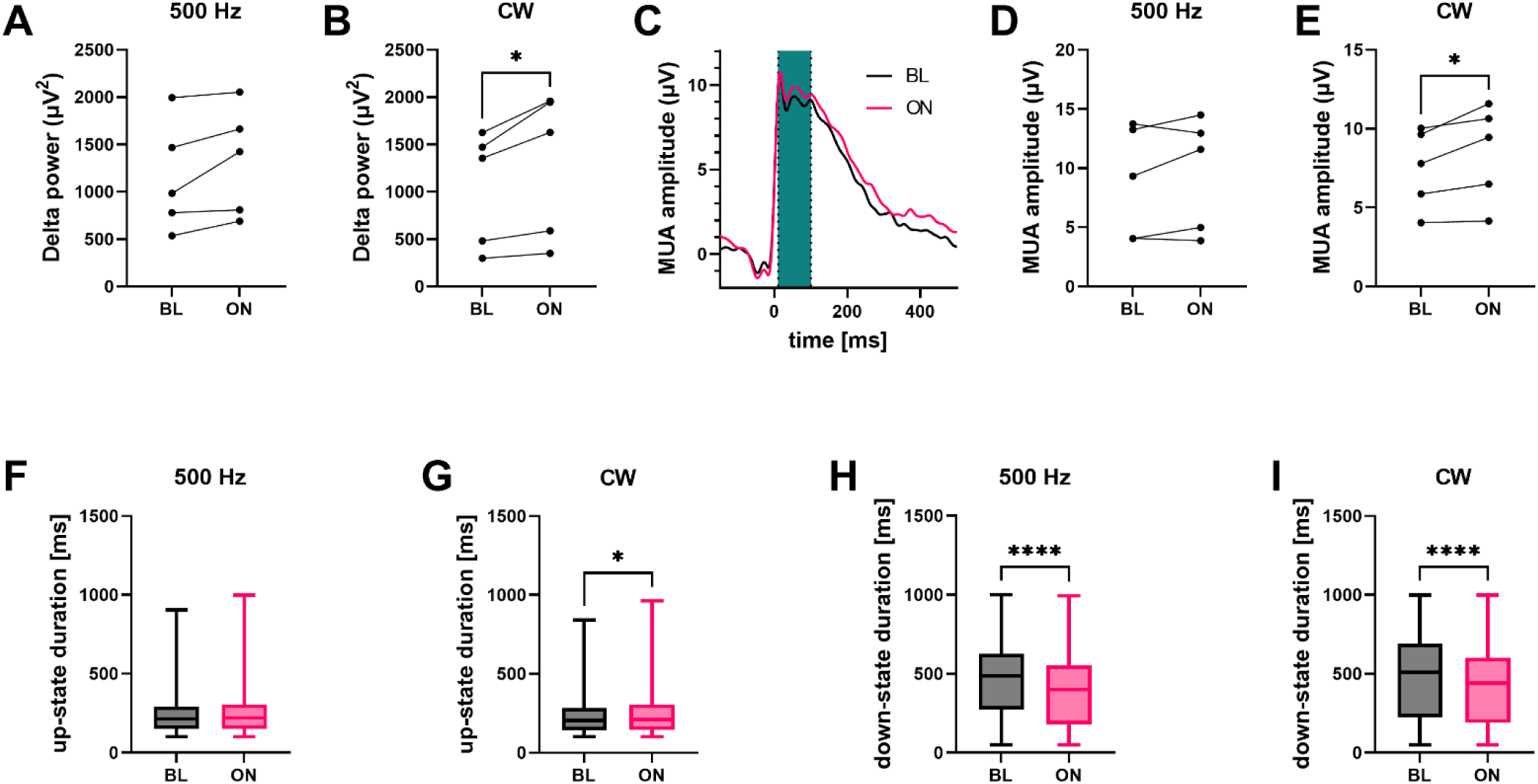
Effect of infrared stimulation on ketamine/xylazine-induced slow-wave activity in wild-type (WT) mice. (A–B) Delta band (0.5–4 Hz) power during baseline (BL) and stimulation (ON) for pulsed (500 Hz; A) and continuous-wave (CW; B) protocols. * p < 0.05. (C) Average cortical multi-unit activity (MUA) aligned to the up-state onsets detected during CW infrared stimulation (pink, ON) and baseline periods (black, OFF) across all WT mice. Up-state onset is at time zero. The green band marks the time window (10 – 100 ms relative to up-state onset) used to calculate MUA amplitudes shown in panels D and E. (D–E) Mean MUA amplitudes within the selected time window during baseline and stimulation for pulsed (D) and CW (E) protocols. * p < 0.05. (F-G) Distribution of the duration of up-states during baseline (BL) and stimulation (ON) periods for pulsed (F) and CW (G) stimulation. Up-states were significantly longer during ON periods for CW stimulation. * p < 0.05. (H-I) Distribution of the duration of down-states during baseline (BL) and stimulation (ON) periods for pulsed (H) and CW (I) stimulation. Down-states were significantly shorter during ON periods compared to baseline. **** p < 0.0001. Boxplots show the median (line), interquartile range (box), and minimum/maximum values (whiskers).

Slow-wave activity consists of two alternating phases: up-states with elevated synaptic activity and increased firing of neurons, and down-states with diminished neuronal firing [48], [55], [54]. Next, we examined average multi-unit activity amplitudes during up-states both for infrared stimulation and baseline periods. Panel C of figure 4 shows the average MUA aligned to up-state onsets from all WT animals during CW stimulation (ON) and related baseline (BL) period. Comparing MUA amplitudes in a 90-ms-long time window after the up-state onset between baseline and ON periods, we found that amplitudes increased during both stimulation modes, with a significant difference observed under CW stimulation (BL vs. ON; 500 Hz: 8.89 µV ± 4.91 µV vs. 9.58 µV ± 4.74 µV, paired t-test, p = 0.2717; Fig. 4D; CW: 7.47 µV ± 2.75 µV vs. 8.46 µV ± 3.16 µV, paired t-test, p = 0.0459). To compare the effects of 500 Hz and CW stimulation, we analyzed changes in MUA amplitudes during ON periods between stimulation modes; however no significant differences were observed (CW vs. 500 Hz; 8.46 µV ± 3.08 µV vs. 9.58 µV ± 4.83 µV, Wilcoxon signed-rank test; p = 0.8125).

We also investigated possible changes in the duration of the up- and down-states (Fig. 4F and G). For up- state durations, no difference was observed during pulsed stimulation (BL vs. ON; 240.40 ms ± 120.34 ms vs. 245.67 ms ± 123.9 ms, Mann-Whitney U test, p = 0.1075; Fig. 4F); however, up-states were significantly longer under CW stimulation (BL vs. ON; 230.59 ms ± 115.63 ms vs. 241.07 ms ± 122.74 ms, Mann-Whitney U test, p = 0.0124; Fig. 4G). Analysis of down-state durations revealed significant differences between baseline and ON periods (Fig. 4H and I). Down-states were significantly shorter during ON periods compared to baseline under both 500 Hz (BL vs. ON; 463.13 ms ± 238.30 ms vs. 385.06 ms ± 225.07 ms, Mann-Whitney U test, p < 0.0001; Fig. 4H) and CW stimulation (BL vs. ON; 481.14 ms ± 267.22 ms vs. 420.42 ms ± 242.88 ms, Mann-Whitney test, p < 0.0001; Fig. 4I).

### 3.2. Comparative effects of infrared neurostimulation in wild-type and TRPV1 knockout mice

Our results demonstrate that infrared neural stimulation significantly modulates cortical activity in anesthetized wild-type mice (Figs. 1-4). To investigate the mechanisms underlying IR stimulation-related responses, particularly the contribution of heat-sensitive ion channels, we performed experiments in TRPV1 knockout (KO) mice (n = 5) applying the same methodology as used in wild-type animals. Because results obtained in wild-type mice were comparable between pulsed and continuous-wave stimulation, with CW stimulation typically producing stronger effects, we focus here on the comparison between WT and KO animals under CW stimulation.

#### 3.2.1 Differences in single unit properties

We first compared the proportion of cortical single units modulated by CW stimulation and their firing rate changes between WT and TRPV1 KO mice (n = 527 units in total; 105.4 ± 50.38 units per mouse; 41.2 ± 10.01 units per recording for CW stimulation). Similarly, as observed in WT mice (Fig. 1B), units in KO animals demonstrated periodic fluctuations in firing rates in response to IR stimulation (Fig. 5A).

**Figure 5.**
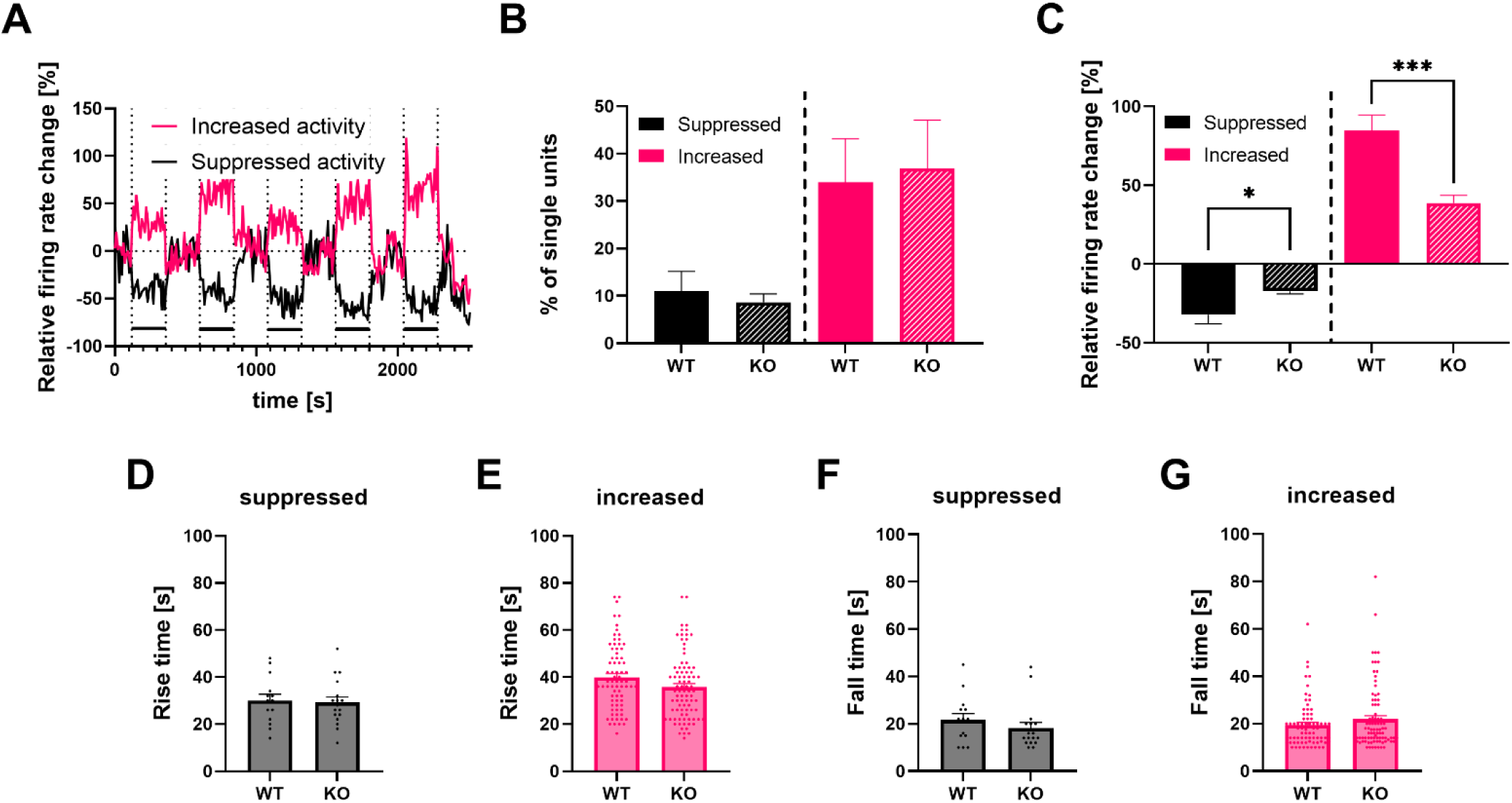
**4** Comparison of the effects of infrared stimulation on somatosensory cortical activity between wild-type (WT) and TRPV1 knockout (KO) mice. (A) Relative firing rate changes of two example single units recorded from a KO mouse. All five stimulation trials are shown. The ON periods are marked with horizontal bars. (B) Proportions of single units with suppressed (black) and increased (red) activity in response to continuous-wave (CW) stimulation in WT and KO animals. (C) Relative firing rate changes of single units from WT and KO animals during CW stimulation. Stimulation-induced changes in firing rates were significantly smaller in KO animals for both types of modulated neurons (suppressed and increased units). * p < 0.05, *** p < 0.001. (D–E) Rise times of single units with suppressed (D) and increased (E) activity.(F–G) Fall times of single units with suppressed (F) and increased (G) activity. Error bars in all panels indicate the standard error of the mean.

A comparison of the proportion of single units with suppressed and increased activity revealed no statistically significant differences between WT and KO animals (WT vs. KO; suppressed: 10.92% ± 9.49% vs. 8.63% ± 3.96%, Mann-Whitney U test, p > 0.9999; increased: 33.95% ± 20.60% vs. 36.89% ± 22.81%, Mann-Whitney U test, p = 0.5476; Fig. 5B). In contrast, analysis of relative firing rate changes revealed significant differences between WT and KO mice. Firing rates of neurons were significantly less suppressed (WT vs. KO; -32.02% ± 21.85% vs. -17.28% ± 7.09%, Student’s t-test, p = 0.0136; Fig. 5C) and less increased (WT vs. KO; 84.57% ± 84.98% vs. 38.84% ± 44.19%, Mann-Whitney U test, p <0.0001; Fig. 5C) in KO animals than in WT mice. These results indicate that CW stimulation exerts a stronger modulatory effect on firing rates in animals with TRPV1 channels.

We next compared the rise and fall times of single units between WT and KO animals (Fig. 5D-G). Rise times of suppressed and increased neurons were not significantly different between KO and WT animals (WT vs. KO; suppressed: 30.00 s ± 10.17 s vs. 29.17 s ± 9.90 s, Mann-Whitney U test, p = 0.7163; Fig. 5D; increased: WT vs. KO; 39.86 s ± 14.20 s vs. 35.72 s ± 13.90 s, Mann-Whitney U test, p = 0.0618; **Error! Reference source not found.**Fig. 5E). Similarly, we found no significant differences in the fall times of neurons between the two animal groups (WT vs. KO; suppressed: 21.64 s ± 10.21 s vs. 18.23 s ± 9.66 s, Mann-Whitney U test, p = 0.2009; increased: 19.37 s ± 9.74 s vs. 21.79 s ± 13.49 s, Mann-Whitney U test, p = 0.3869; Fig. 5F and G).

Furthermore, we examined neuron-type-specific differences between WT and KO animals. In KO animals, the proportion of wide-spiking interneurons was somewhat higher (WT vs. KO; ∼9% vs. ∼24%; Fig. 2C vs. 6A; Supplementary Table S6) and the proportion of narrow-spiking interneurons slightly lower compared to WT animals (WT vs. KO; ∼34% vs. ∼24%; Fig. 2C vs. 6A; Supplementary Table S6), while the proportion of principal cells was similar (WT vs. KO; ∼57% vs. ∼52%; Fig. 2C vs. 6A; Supplementary Table S6). While there was no significant difference across genotypes in the proportion of both interneuron types with suppressed or increased activity (WT vs. KO; N-I increased: 36.03% ± 21.24% vs. 39.89% ± 9.13%, Mann-Whitney U test, p>0.9999; N-I suppressed: 1.32% ± 1.81% vs. 10.46% ± 6.03%, Mann-Whitney U test, p = 0.0635; W-I increased: 48.24% ± 29.65% vs. 33.74% ± 7.97%, Mann-Whitney U test, p = 0.1508; W-I suppressed: 6.48% ± 6.13% vs. 2.59% ± 3.63%, Mann-Whitney U test, p = 0.2857; Fig. 6B and C), significantly less excitatory principal cells were found with suppressed and more with increased activity in KO animals (WT vs. KO; suppressed: 15.29% ± 10.13% vs. 6.65% ± 1.69%, Mann–Whitney U test, p = 0.0079; increased: 42.46% ± 11.09% vs. 65.48% ± 8.54%, Mann–Whitney U test, p = 0.0159; Fig. 6D).

**Figure 6.**
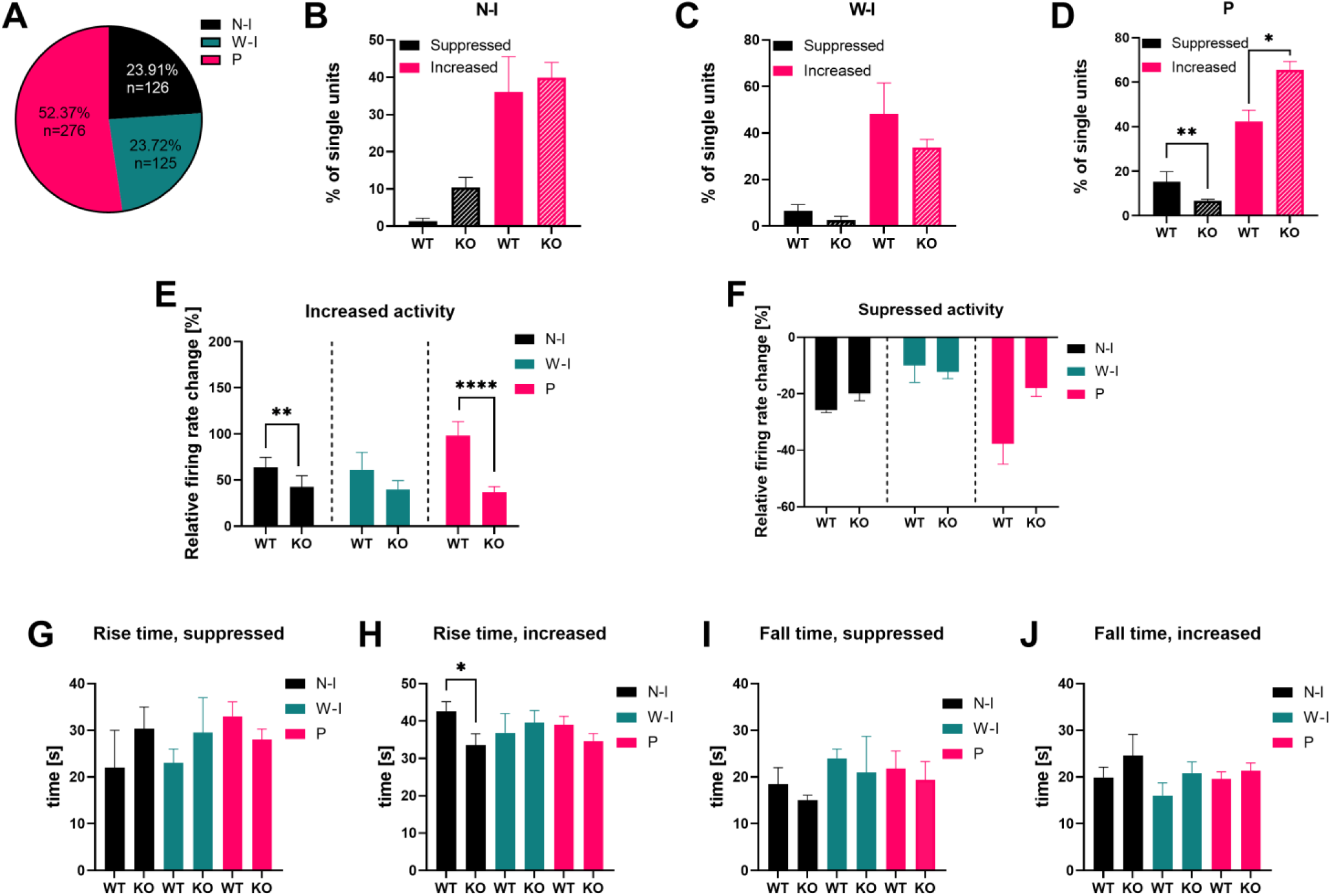
Neuron type–specific differences between wild-type (WT) and TRPV1 knockout (KO) mice to continuous-wave (CW) stimulation. (A) Proportions of the three neuron types (n = 527 single units in total). Percentages and cell counts are indicated for each type. P, principal cells; N-I, narrow-spiking interneurons; W-I, wide-spiking interneurons. (B–D) Proportions of single units with suppressed and increased activity in WT and KO mice for the three neuron types. In KO animals, significantly fewer principal cells were suppressed, and significantly more showed increased activity to CW stimulation.* p < 0.05, ** p < 0.01. (E) Relative firing rate changes of the three cell types with increased activity. In KO animals, the effect of CW stimulation on firing rates was significantly weaker for narrow-firing interneurons and principal cells. ** p < 0.01, **** p < 0.0001. (F) Relative firing rate changes of the three neuron types with suppressed activity to CW stimulation. (G–H) Rise times of the three neuron types with suppressed (G) and increased (H) activity. * p < 0.05. (I-J) Fall times of interneurons and principal cells with suppressed (I) and increased (J) activity. Error bars in all panels represent the standard error of the mean.

Analysis of relative firing rate changes in single units with increased activity revealed that KO animals exhibited lower firing rate changes for all three cell types (Fig. 6E), and these differences were significant for narrow-spiking interneurons and principal cells (WT vs. KO; N-I: 63.66% ± 49.16% vs. 42.40% ± 52.81%, Mann–Whitney U test, p = 0.0472; W-I: 61.09% ± 49.82% vs. 39.98% ± 44.20%, Mann–Whitney U test, p = 0.3490; P: 98.28% ± 99.80% vs. 36.98% ± 40.99%, Mann–Whitney U test, p < 0.0001; Fig. 6E). A similar trend was observed in suppressed cells, although the differences were not statistically significant (WT vs. KO; N-I: -26.82% ± 1.20% vs. -19.90% ± 6.26%, Mann–Whitney U test, p = 0.1429; W-I: -9.94% ± 8.58% vs. -12.25% ± 4.78%, Mann–Whitney U test, p = 0.5333; P: -37.67% ± 22.99% vs. -17.90% ± 7.96%, Mann–Whitney U test, p = 0.0553; Fig. 6F). Rise and fall times of different neuron types were similar between WT and KO animals, except for narrow-spiking interneurons with increased activity, where rise times were significantly faster in KO mice (WT vs. KO; 42.64 s ± 11.88 s vs. 33.85 s ± 13.12 s, Mann-Whitney U test, p = 0.0366; Fig. 6H; Supplementary Table S7).

We next compared layer-dependent effects of CW stimulation in WT and KO animals across the superficial/input layers (L1/2/3/4; 19.4 ± 9.1 units per KO mouse, n = 97 units in total), layer 5 (L5; 42.4 ± 23.9 units per KO mouse, n = 212 units in total), and layer 6 (L6; 43.6 ± 36.7 units per KO mouse, n = 218 units in total; Supplementary Table S8). A close examination of the proportion of single units revealed no significant differences across layers, although neurons exhibiting increased activity were found to be more prevalentin layers 1-4 and layer 5 of KO animals (Fig. 7A-C; Supplementary Table S9).

**Figure 7.**
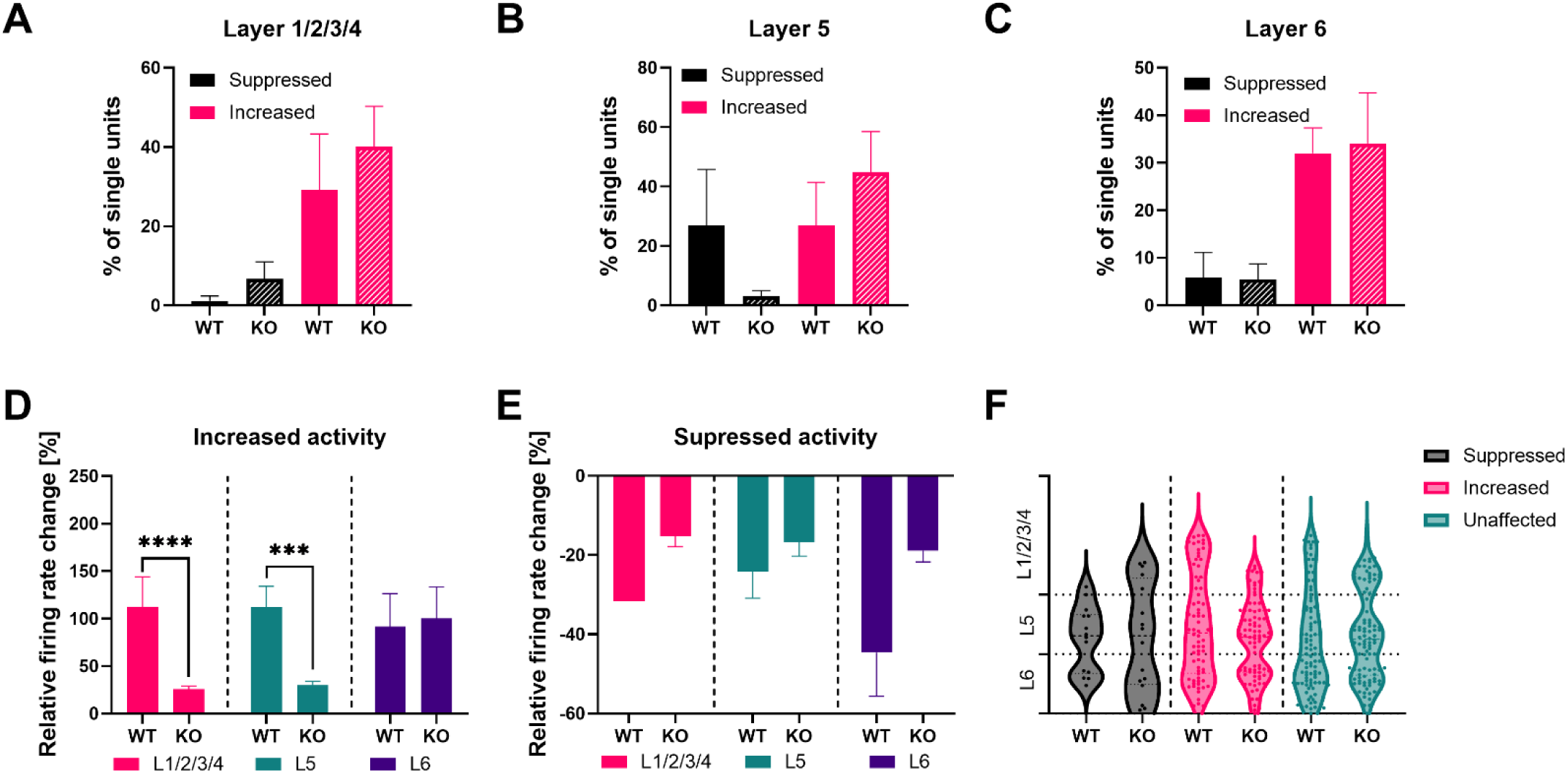
5 Layer-specific differences between wild-type (WT) and TRPV1 knockout (KO) mice to continuous-wave (CW) stimulation. (A-C) Percentages of single units exhibiting suppressed or increased activity in superficial/input layers (L1/2/3/4, A), layer 5 (L5, B) and layer 6 (L6, C) in WT and KO animals. (D-E) Relative firing rate changes in neurons with increased (D) and suppressed (E) activity across all three layer groups. Firing rate changes in KO animals to CW stimulation were significantly lower in the superficial/input layers and layer 5 compared to WT animals. *** p < 0.001, **** p < 0.0001. (F) Distribution of single units with suppressed, increased, or unaffected responses to CW stimulation, plotted by cortical depth. Each point represents a single unit. The y-axis indicates the normalized depth of the neurons, with dashed horizontal lines marking the boundaries between layers. Error bars in all panels show standard error of the mean.

Upon examination of the relative firing rate changes of single units exhibiting increased activity, KO animals demonstrated significantly lower firing rate changes in comparison to WT mice within the superficial/input layers (WT vs. KO; 112.14% ± 152.08% vs. 25.66% ± 12.30%, Mann-Whitney U test, p < 0.0001; Fig. 7D) and layer 5 (WT vs. KO; 111.74% ± 112.07% vs. 30.02% ± 24.65%, Mann-Whitney U test, p = 0.0003; Fig. 7D), while increase of firing rates was comparable in layer 6 (WT vs. KO; 91.47% ± 178.04% vs. 100.14% ± 193.39%, Mann-Whitney U test, p = 0.6948; Fig. 7D). In the case of suppressed cells, the mean firing rate change of KO animals was consistently lower across all layers compared to WT mice but no significant difference was found (WT vs. KO; L1/2/3/4: -31.64% ± 0.00% vs. -15.20% ± 6.04%, Student’s t-test, p = 0.0678, L5: -24.19% ± 18.87% vs. -16.67% ± 7.22%, Mann-Whitney U test, p = 0.4606, L6: -44.63% ± 24.64% vs. -18.89% ± 7.97%, Mann-Whitney U test, p = 0.0932; Fig. 7E). As illustrated in panel F of figure 7 showing the distribution of increased, suppressed and unaffected units plotted against the normalized cortical depth, in wild-type animals, suppressed activity was primarily localized to layers 5 and 6. In contrast, KO animals showed a broader distribution of suppressed activity, including layer 6 as well as the superficial and input layers. For increased activity, neurons were mainly found in layer 6 and the superficial and input layers for WT mice, while KO animals showed a higher proportion of activated cells in layers 5 and 6.

Our anatomical analysis in WT mice revealed that TRPV1 is expressed across all cortical layers of the somatosensory cortex in both excitatory and inhibitory neurons (Fig. 8), with TRPV1 mRNA expression in layer 6 restricted to glutamatergic neurons (Fig. 8F). This distribution aligns with our electrophysiological findings, which showed that no single cortical layer plays a dominant role in IR stimulation–mediated changes (Fig. 3). In TRPV1-deficient mice, IR stimulation produced a weaker effect on firing rates (Fig. 5C); however, laminar analysis indicated that this reduction was absent in layer 6 units with increased activity, consistent with the anatomical observation that TRPV1 expression in layer 6 differs from other layers (Fig. 7D).

**Figure 8.**
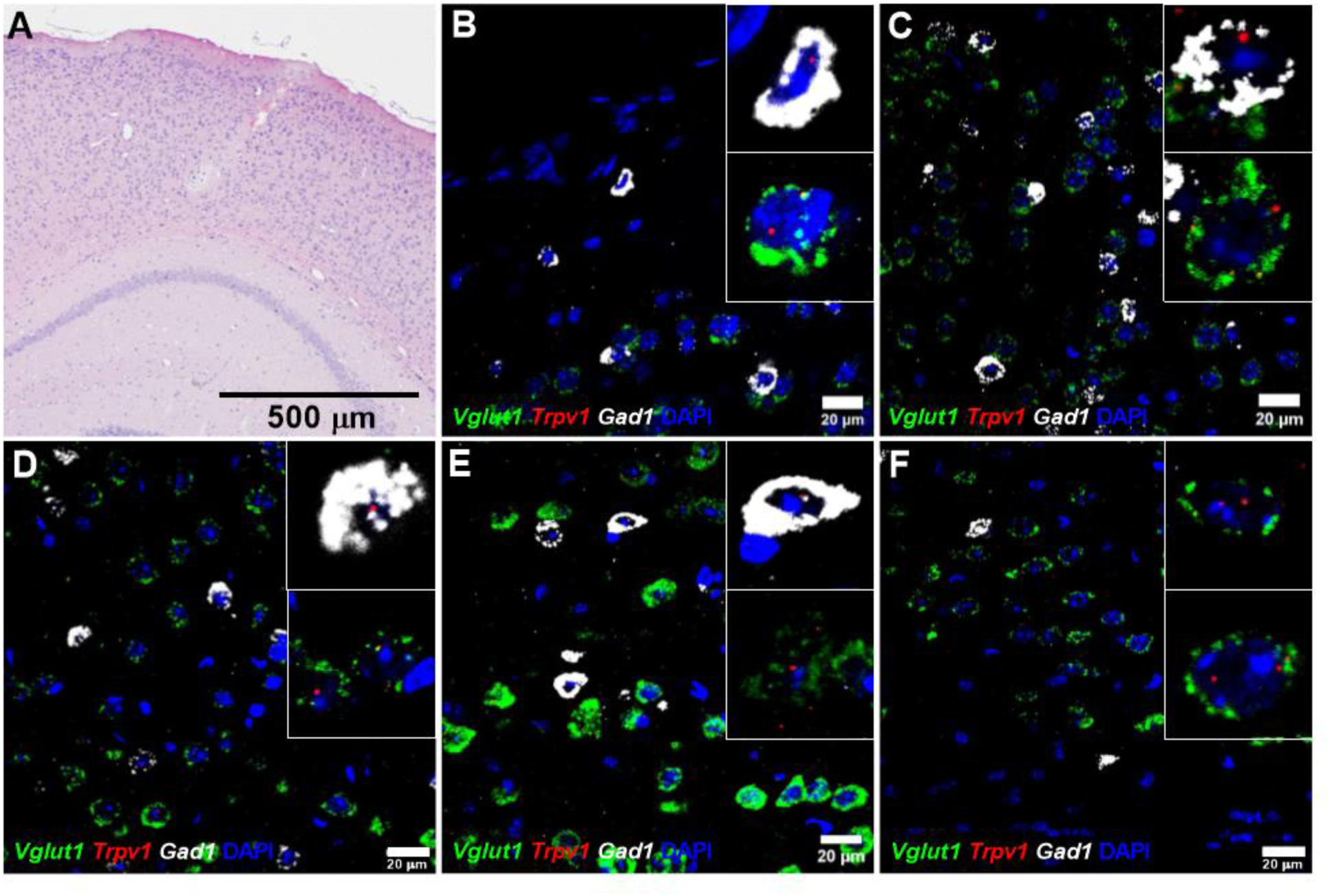
TRPV1 mRNA expression in the primary somatosensory cortex of wild-type mice. (A) Hematoxilin-eosin staining of the somatosensory cortex from a treated animal. (B-F) Representative images showing mRNA expression across cortical layers. Colocalization of the Trpv1 mRNA dots (red) with Vglut1 (green; excitatory neurons) and Gad1 (white; inhibitory neurons) is visible in layer 1 (B), layer 2/3 (C), layer 4 (D), and layer 5 (E). In layer 6 (F), expression was restricted to excitatory neurons. Nuclei are counterstained with DAPI (blue). Scale bars: 500 µm (A) and 20 µm (B-F).

#### 3.2.2 Differences in the effects on ketamine/xylazine-induced slow-wave activity

We compared CW stimulation-induced changes in the spectral power across all investigated frequency bands relative to baseline between WT and KO mice (Supplementary Fig. 4). While the change in delta and theta power was slightly larger in KO animals, differences were not significant (Supplementary Table S10). Change of power in alpha, beta and gamma bands were low and similar between WT and KO mice, without any significant differences (Supplementary Fig. 4). We next evaluated stimulation-related changes in cortical MUA amplitudes during up-states (measured from 10 to 100 ms after up-state onset) in WT and KO animals (Fig. 9A). Relative changes in MUA amplitudes were not significantly different and were slightly lower in KO animals suggesting a weaker stimulation effect (WT vs. KO; 12.17% ± 8.24% vs. 10.99% ± 15.61%, Mann-Whitney U test, p = 0.8413; Fig. 9A). However, it is important to note that baseline MUA amplitudes were significantly different between WT and KO mice (WT vs. KO; 7.47 µV ± 2.75 µV vs. 10.82 µV ± 3.77 µV, Mann-Whitney U test, p = 0.0030).

**Figure 9.**
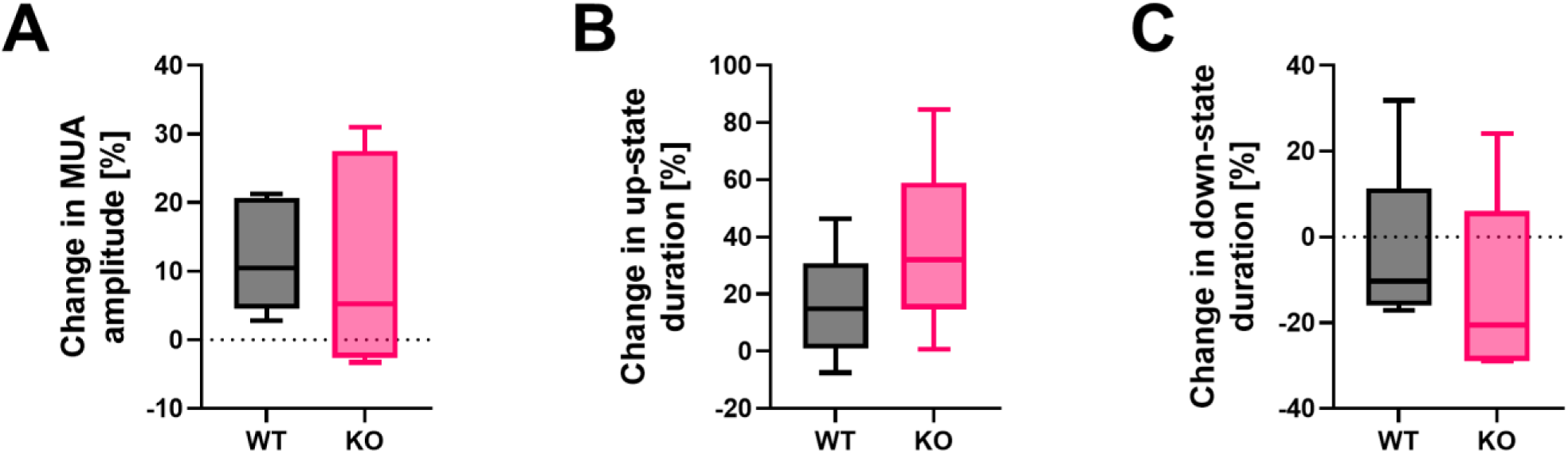
6 Comparison of the effects of infrared stimulation on slow-wave activity in wild-type (WT) and TRPV1 knockout (KO) mice. (A) Changes in up-state-locked multi-unit activity (MUA) amplitudes to continuous-wave (CW) stimulation relative to baseline amplitudes in WT and KO mice. (B-C) CW stimulation-induced changes in up-state (B) and down-state (C) durations in WT and KO animals relative to baseline state durations. Boxplots show the median (line), interquartile range (box), and minimum/maximum values (whiskers). No significant differences were detected between WT and KO mice.

Finally, we investigated changes in up- and down-state durations to stimulation (Fig. 9B and C). Up-states became longer also in KO animals showing a slightly larger increase in state duration; however, the difference between WT and KO mice was not significant (WT vs. KO; 15.54% ± 19.53% vs. 35.83% ± 30.37%, Mann-Whitney U test, p = 0.3095; Fig. 9B). Similarly, down-states became shorter in KO animals but the difference between WT and KO mice was not significant (WT vs. KO; -3.93% ± 20.27% vs. -13.27% ± 22.04%, Mann-Whitney U test, p = 0.2222; Fig. 9C). Baseline up- and down-state durations were significantly different between WT and KO mice, with longer up-states and shorter down-states in KO animals (WT vs. KO; up-state: 230.58 ms ± 105.62 ms vs. 359.04 ms ±146.09 ms, Mann-Whitney U test, p < 0.0001; down-state: 529.56 ms ± 319.38 ms vs. 506.02 ms ± 226.35 ms, Mann-Whitney U test, p = 0.0049).

## 4. Discussion

Understanding how infrared neural stimulation modulates neuronal excitability has been a growing area of interest over the past 15 years. In the initial stages of research, the focus was predominantly on *in vitro* and *ex vivo* models. The objective of these models was to reveal the underlying excitatory and inhibitory mechanisms of IR stimulation at the cellular level. While these models have provided valuable mechanistic insights, *in vivo* investigations have been comparatively rare and have primarily relied on indirect measurements such as BOLD functional imaging [28], [56], two-photon calcium imaging [6], [57], or low-density extracellular recordings [58]. While these approaches have been demonstrated to be effective in the assessment of bulk tissue responses, they frequently lack the requisite resolution to capture single-cell activity patterns and layer-specific dynamics within an intact cortical environment.

This study addresses this gap by using high-density laminar microelectrode recordings to simultaneously monitor neuronal spiking across cortical layers in the mouse somatosensory cortex during INS. Furthermore, by using TRVP1-deficient mice, we investigated the role of TRPV1 channels in mediating IR-evoked responses. To our knowledge, this is the first *in vivo* study to systematically evaluate the effects of INS on cortical activity in both TRPV1 KO and WT mouse models.

### 4.1. Effects of infrared stimulation on somatosensory activity in anesthetized wild-type mice

Local IR stimulation in wild-type mice induced significant changes in neuronal firing near the recording probe, with a large proportion of neurons increasing their firing rate and a smaller fraction decreasing activity. The proportion of responsive neurons, along with firing rate changes and rise/fall times, was comparable to our previous observations in the somatosensory cortex of anesthetized rats [37]. Furthermore, no notable differences were detected across neuronal subtypes or cortical layers in the examined single unit properties, suggesting that both interneurons and principal cells, as well as all cortical layers, contribute to IR-stimulation-related responses. Moreover, infrared stimulation strongly affected slow-wave activity (0.5 - 4 Hz) induced by ketamine/xylazine, including a significant increase in delta power and in MUA amplitudes during up-states, consistent with our earlier findings in anesthetized rats [54]. However, the effects of IR stimulation on up- and down-state durations were opposite in mice and rats: in mice, up-states were prolonged, and down-states shortened, whereas in rats, the reverse pattern was observed [54]. In addition, our results indicate that continuous-wave stimulation produced stronger effects compared to 500 Hz stimulation, reflected in larger changes in neuronal firing rates, lower rise times, greater increase in delta power and MUA amplitudes, and longer up-state durations.

Infrared stimulation modulates slow-wave activity - by altering LFP and MUA amplitudes, up- and down-state durations, and population spiking synchrony - likely through localized tissue heating. These effects are consistent with prior findings linking warmer tissue to increased low-frequency LFP power and more synchronized neuronal activity [54]. Previous research has highlighted the strong influence of cortical temperature on both slow and fast cortical rhythms [59], [60], [61]. For example, Reig and colleagues [59] showed in visual cortical slices that higher temperatures increased low-frequency LFP power and sharpened population activity during slow-wave activity. Kalmbach and Waters [60] demonstrated that exposed cortical surfaces in cranial windows are cooler than core brain temperature, leading to altered slow-wave dynamics, with warmer tissue associated with shorter up-states. Similarly, Sheroziya and Timofeev [61] found that moderate cooling of the somatosensory cortex eliminated down-states during slow oscillations, indicating that even small temperature changes can substantially modulate network excitability and state transitions. The effects of IR stimulation may also vary across cortical areas: slow and delta oscillations were enhanced in the parietal association cortex but reduced in the primary somatosensory cortex of anesthetized rats [54]. This regional variability likely reflects differences in laminar structure, cortical thickness, baseline tissue temperature, and fiber placement.

Astrocytes may also contribute to IR-induced modulation of slow waves. Calcium imaging studies show that pulsed infrared light evokes slow astrocytic responses, influencing the generation and synchronization of cortical slow waves [62], [63]. Thus, IR-induced tissue heating likely affects both neurons and astrocytes, shaping the population-level cortical dynamics observed in LFP and MUA measurements.

Layer-specific analysis showed slight, but not statistically significant, stimulation-related differences between layers. For example, stronger suppression was observed in deeper layers, whereas superficial and input layers showed a higher proportion of neurons with increased activity. Although the IR fiber was positioned approximately 600 μm below the cortical surface to primarily target layer 5, thermal effects extend over ∼1 mm, and neural activity rapidly propagates across layers under anesthesia [34], [37], [61]. Consequently, initially layer-specific effects may become less distinct during stimulation, which could explain the considerable variability in cortical responses observed here and in previous studies [34], [37]. A key limitation, therefore, is that the applied IR stimulation may lack the spatial precision required to selectively manipulate individual cortical layers. Alternatively, the absence of strict layer-specific responses might reflect the intrinsic interconnectivity of cortical microcircuits, where perturbations in one layer are rapidly integrated across the network.

### 4.2. TRPV1 as a mediator of infrared stimulation

In TRPV1-deficient animals, the proportion of neurons responding to IR stimulation was similar to that in WT mice. However, firing rate changes were significantly smaller in both suppressed and activated neuron populations, indicating a weaker response to IR stimulation in KO mice. This reduced effect was consistent across cell types and cortical layers with the exception of layer 6. In contrast, the impact of infrared stimulation on slow-wave dynamics was similar between KO and WT mice, with no significant differences observed in the properties examined.

As noted above, changes in firing rates were genotype-dependent: modulated neurons in KO animals showed smaller firing rate changes compared to WT animals, indicating that TRPV1 contributes to both excitatory and inhibitory responses. At the network level, KO animals exhibited slightly larger, but not statistically significant changes in delta band power under ketamine/xylazine anesthesia, as well as greater changes in up- and down-state durations compared to WT mice. However, baseline values differed significantly between KO and WT animals, suggesting that TRPV1 may also influence spontaneous slow-wave dynamics.

We found that TRPV1 is expressed throughout all layers of the somatosensory cortex in both excitatory and inhibitory neurons, except in layer 6, where its expression is confined to glutamatergic cells. This widespread distribution aligns with our electrophysiological data, which show that IR-induced responses are not dominated by any single layer. In TRPV1 knockout animals, IR-driven modulation of neuronal activity is reduced in layers 1–5, but for neurons with increased activity in layer 6, responses are comparable to those in WT animals, indicating that TRPV1 partially mediates layer-specific excitability.

At the molecular level, thermosensitive TRP channels play a central role in mediating infrared responses, with TRPV1 acting as the primary contributor [24], [63], [64]. Knockout studies and pharmacological inhibition consistently demonstrate that disrupting TRPV1 significantly reduces IR-evoked neuronal activity [24], [63], [64]. Other heat-sensitive TRP channels, such as TRPV2 and TRPV3, are less likely to contribute under typical IR stimulation due to their higher activation thresholds [24][34]. Nonetheless, TRPV1 is not the sole mediator, as IR effects on neuronal activity persist in TRPV1 KO animals, as shown here. Additional thermosensitive channels, including TRPV2–4, as well as mechanosensitive channels such as TREK, may support infrared-induced modulation of cortical activity. Considering that cortical temperatures during stimulation likely remain below ∼40 °C, channels with lower activation thresholds, such as TRPV3 (>32 °C), TRPV4 (>27 °C), TRPM2, and mechanosensitive channels including Piezo and TREK/TRAAK, may contribute to the residual activity [24], [34], [64], [65]. Moreover, IR stimulation can trigger calcium signaling in astrocytes, suggesting that neuroglial interactions also shape the overall network response [63]. Collectively, these findings highlight a multifaceted mechanism in which TRPV1 serves as an important molecular link between infrared-induced heating and neuronal excitability, while other channels and glial signaling may modulate and fine-tune the response.

### 4.3. Limitations

Our study has a few limitations that should be considered when interpreting the results. Experiments were performed under ketamine/xylazine anesthesia, which alters physiological cortical network activity and may have influenced the observed effects of stimulation. Furthermore, we only assessed IR stimulation-related responses in a single brain area. Previous work indicates that infrared stimulation can produce region-specific effects; for instance, enhancing slow and delta oscillations in the parietal association cortex but reducing them in the somatosensory cortex [54]. These findings suggest that IR-induced responses may depend on both local circuitry and cortical area, highlighting the need for more systematic investigations.

Although fiber placement was carefully controlled, IR stimulation-related effects might differ if the fiber were positioned in another layer, such as the supragranular layers. Slow-wave activity is primarily generated in layer 5, so stimulating in a different layer could potentially produce distinct responses. However, this effect may be limited, as IR stimulation affects an area of roughly 1 mm which is comparable to the thickness of the mouse cortex. In addition, we compared only two stimulation modes (continuous-wave and 500 Hz pulsed); exploring additional illumination patterns and stimulation frequencies could provide a more comprehensive understanding of infrared neuromodulation dynamics. Finally, it is worth noting that mRNA expression does not always correlate perfectly with protein expression. Nevertheless, our RNAscope experiments allowed us to infer the presence of TRPV1 channels, supporting the molecular basis for the observed effects.

## Acknowledgments

This work is dedicated to the memory of our coauthor Zoltán Fekete (deceased December 2024), whose contributions were essential to this study. The research leading to these results has received funding from the Hungarian Brain Research Program Grant (NAP2022-I-8/2022, NAP2022-I-2/2022). Project no. 150574 has been implemented with the support provided by the Ministry of Culture and Innovation of Hungary from the National Research, Development, and Innovation Fund, financed under the STARTING_24 funding scheme. R. F. was supported by the Bolyai János Scholarship of the Hungarian Academy of Sciences. Zs. B-L. and Á. Cs. H. were supported by the EKÖP-24-4 University Research Scholarship Program (EKÖP-24-3-II-PPKE-84 and EKÖP-24-4-II-PPKE-59, respectively) of Ministry for Culture and Innovation of Hungary from the National Fund for Research. The research was performed in collaboration with ‘’István Ábrahám’’ Nano-Bio-Imaging Core Facility, University of Pécs.

## Author contributions

**Zsófia Balogh-Lantos:** Data curation, Formal analysis, Methodology, Software, Validation, Visualization, Writing— original draft, Writing—review & editing. **Ágoston Csaba Horváth:** Data curation, Investigation, Writing—review & editing. **Katalin Rozmer:** Investigation, Methodology, Visualization, Writing—review & editing. **Richárd Fiáth:** Conceptualization, Data curation, Investigation, Methodology, Resources, Supervision, Funding acquisition, Writing— original draft, Writing—review & editing. **Zsuzsanna Helyes:** Conceptualization, Funding acquisition, Resources, Supervision, Writing—review & editing. **Zoltán Fekete:** Conceptualization, Methodology, Project administration, Supervision, Funding acquisition, Writing— original draft.

## Data availability

Data will be provided upon reasonable request sent to the corresponding author.

