## Supplementary Material for "Focal infrared stimulation modulates somatosensory cortical activity in mice: evidence for TRPV1 ion channel involvement"

Zsófia Balogh-Lantos^a,b^, Ágoston Csaba Horváth^b^, Katalin Rozmer ^c,d,e,f^, Richárd Fiáth^b,g,^**^*^**, Zsuzsanna Helyes^c,d,e,1^, Zoltán Fekete^b,g,1^**^,†^**

^a^ Roska Tamás Doctoral School of Sciences and Technology, Faculty of Information Technology and Bionics, Pázmány Péter Catholic University, Práter utca 50/A, 1083 Budapest, Hungary

^b^ Research Group for Implantable Microsystems, Faculty of Information Technology and Bionics, Pázmány Péter Catholic University, Práter utca 50/A, 1083 Budapest, Hungary

^c^ Department of Pharmacology and Pharmacotherapy, Medical School, University of Pécs, Szigeti út 12., 7624 Pécs, Hungary

^d^ HUN-REN-PTE Chronic Pain research Group, Szigeti út 12., 7624 Pécs, Hungary

^e^ National Laboratory for Drug Research and Development, Magyar tudósok körútja 2., 1117 Budapest, Hungary

^f^ Institute of Pharmaceutical Chemistry, Faculty of Pharmacy, University of Pécs, Rókus utca 4, 7624 Pécs, Hungary

^g^ Institute of Cognitive Neuroscience and Psychology, HUN-REN Research Centre for Natural Sciences, Magyar tudósok körútja 2., 1117 Budapest, Hungary

**^1^** These authors contributed equally to this work.

**Supplementary Figures**

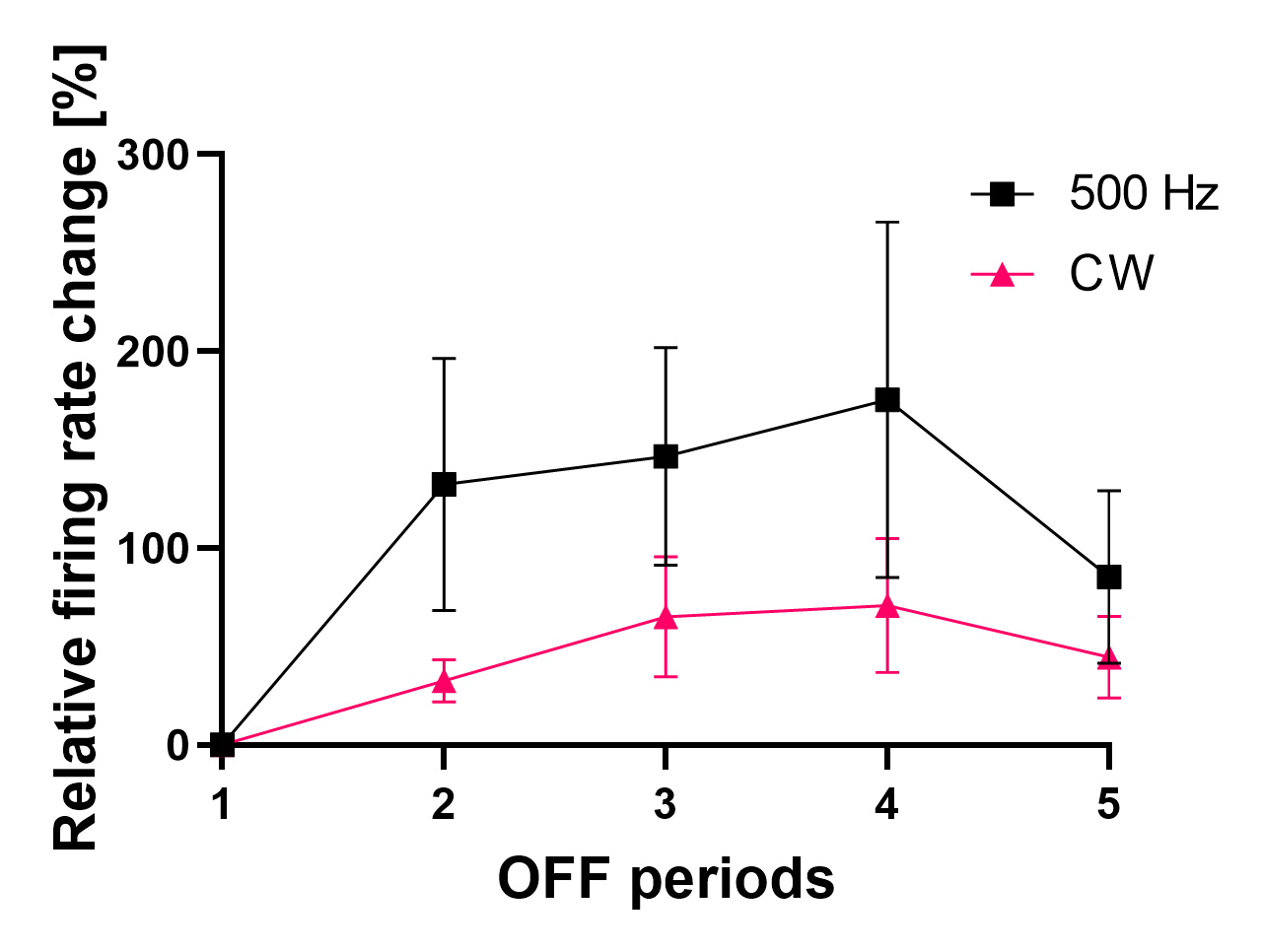

**Supplementary Figure S1.** Firing rate changes of single units in wild-type mice (n = 5) during recovery (OFF) periods preceding infrared stimulation, shown relative to the first baseline period, for both pulsed (500 Hz) and continuous-wave (CW) stimulation modes. Error bars indicate the standard error of the mean.

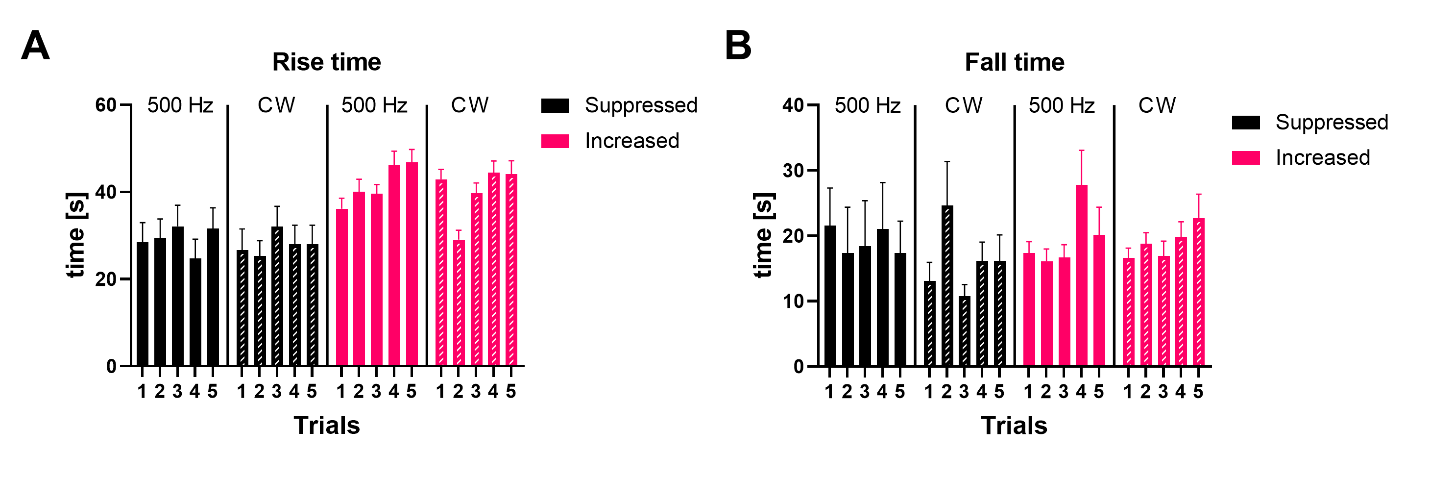
**Supplementary Figure S2.** Trial-to-trial variability of rise (A) and fall (B) times of single units showing suppressed (black) or increased (red) activity in wild-type mice (n = 5) during pulsed (500 Hz) and continuous-wave (CW) stimulation. Error bars in all panels indicate the standard error of the mean. Rise and fall times did not differ significantly across trials.

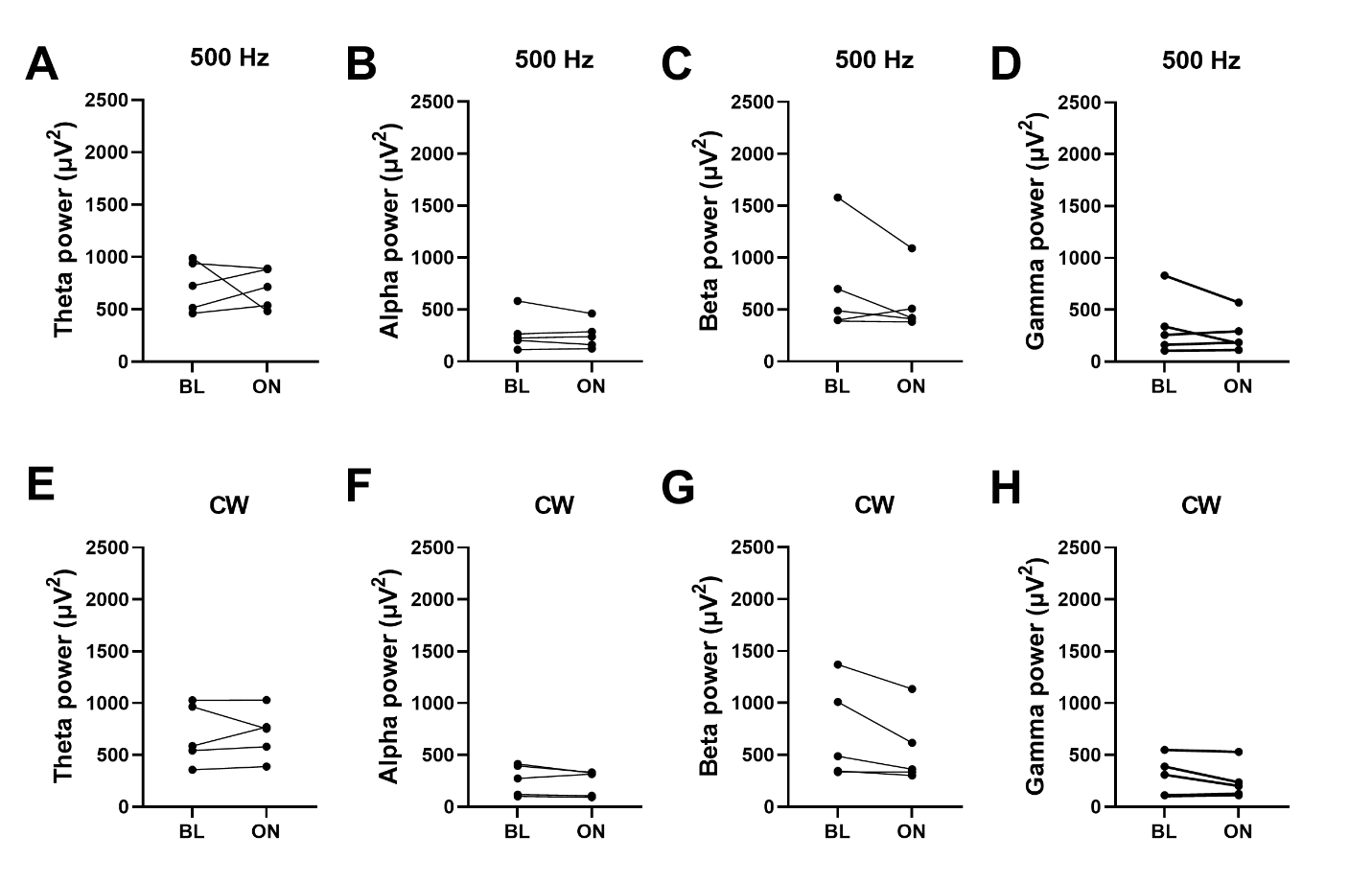
**Supplementary Figure S3.** Average spectral power in the theta (4-8 Hz; A, E), alpha (8-12 Hz; B, F), beta (12-30 Hz; C, G) and gamma (30-100 Hz; D, H) bands during baseline (BL) and stimulation (ON) periods in wild-type mice (n = 5). Data are shown for pulsed (500 Hz; A-D) and continuous-wave (CW; E-H) stimulation. No significant differences in spectral power were observed between baseline and stimulation for any frequency band.

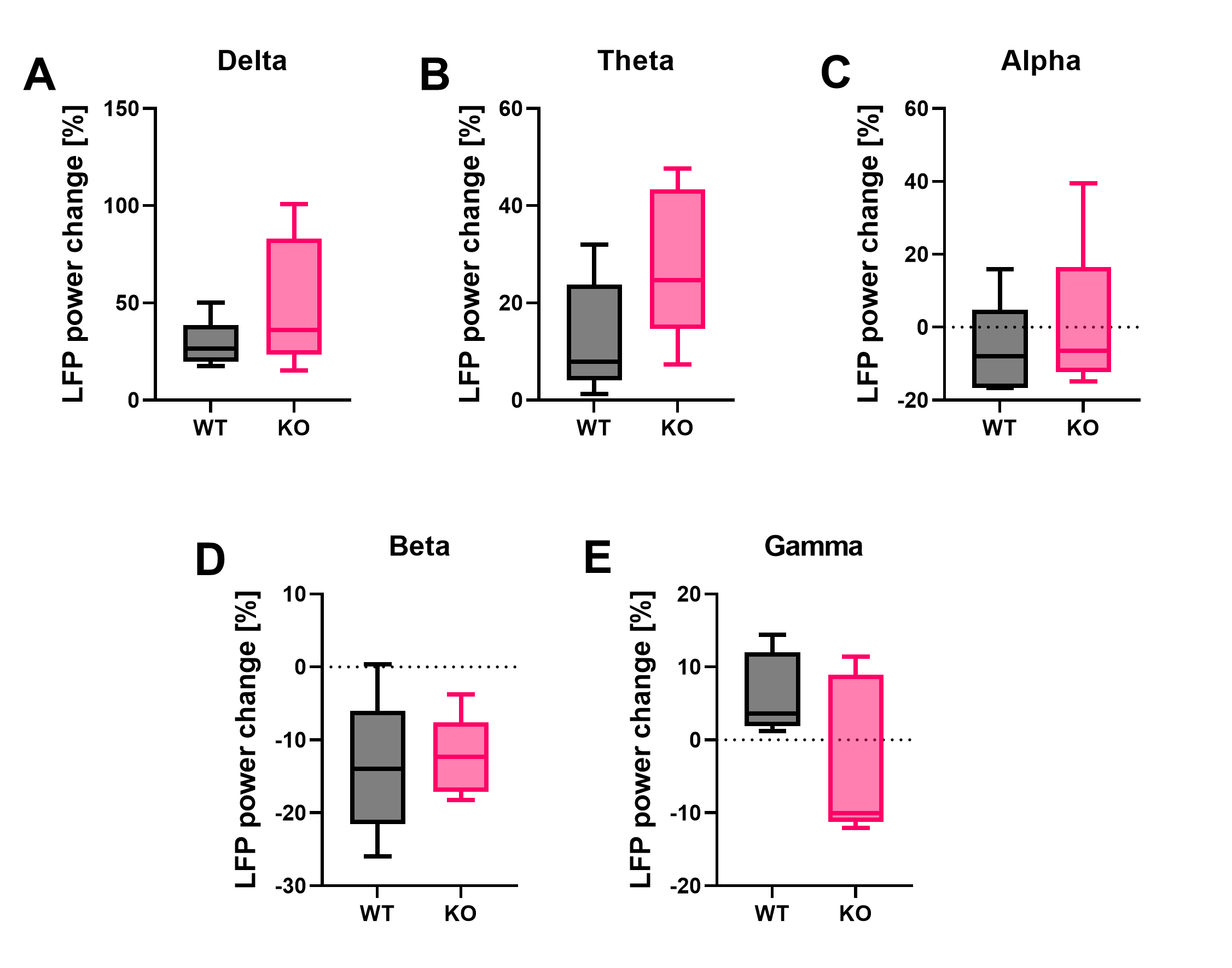
**Supplementary Figure S4.** Continuous-wave infrared stimulation-induced changes in spectral power of the delta (0.5-4 Hz; A), theta (4-8 Hz; B), alpha (8-12 Hz; C), beta (12-30 Hz; D), and gamma (30-100 Hz; E) bands relative to baseline periods for wild-type (WT; n = 5) and TRPV1 knockout (KO; n = 5) mice. Boxplots show the median (line), interquartile range (box), and minimum/maximum values (whiskers). No significant differences in spectral power were observed between WT and KO mice across frequency bands.

**Supplementary Tables**

|  | **Narrow-spiking interneurons [%]** | **Wide-spiking interneurons [%]** | **Principal cells**  **[%]** |
| --- | --- | --- | --- |
| **WT Mouse 1** | 25.00 | 0.00 | 75.00 |
| **WT Mouse 2** | 40.91 | 11.36 | 47.73 |
| **WT Mouse 3** | 21.95 | 24.39 | 53.66 |
| **WT Mouse 4** | 42.86 | 8.93 | 48.21 |
| **WT Mouse 5** | 31.52 | 5.45 | 63.03 |

**Supplementary Table S1.** Percentages of narrow-spiking interneurons, wide-spiking interneurons, and principal cells identified in each wild-type (WT) mouse.

|  | **Layers 1-4** | **Layer 5** | **Layer 6** |
| --- | --- | --- | --- |
| **WT Mouse 1** | 2 | 0 | 18 |
| **WT Mouse 2** | 40 | 25 | 23 |
| **WT Mouse 3** | 2 | 20 | 19 |
| **WT Mouse 4** | 4 | 8 | 44 |
| **WT Mouse 5** | 28 | 60 | 77 |
| **Total** | 76 | 113 | 181 |

**Supplementary Table S2.** Number of single units detected in each cortical layer group for each wild-type (WT) mouse.

|  |  | **Layers 1-4** | **Layer 5** | **Layer 6** | **Statistical test** | **p-value** |
| --- | --- | --- | --- | --- | --- | --- |
| Suppressed | 500 Hz | 7.27% ± 16.26% | 4.44% ± 9.94% | 5.06% ± 4.84% | Friedman | 0.7778 |
|  | CW | 1.18% ± 2.63% | 27.12% ± 41.87% | 6.24% ± 12.54% | Friedman | 0.4074 |
| Increased | 500 Hz | 13.96% ± 15.88% | 33.98% ± 23.25% | 44.82% ± 22.34% | Friedman | 0.0293 |
|  | CW | 28.87% ± 31.24% | 27.20% ± 32.35% | 30.94% ± 13.38% | Friedman | 0.5818 |

**Supplementary Table S3.** Proportions of single units with suppressed or increased activity across cortical layers during pulsed (500 Hz) and continuous-wave (CW) stimulation in wild-type mice (mean ± standard deviation). Statistical comparisons between layers and conditions are reported, including the statistical tests used and corresponding p-values. Although the Friedman test indicated a significant difference between the proportions of units with increased activity for 500 Hz stimulation, no significant differences were found in post-hoc pairwise comparisons.

|  |  | **Layers 1-4** | **Layer 5** | **Layer 6** | **Statistical test** | **p-value** |
| --- | --- | --- | --- | --- | --- | --- |
| Suppressed | 500 Hz | -24.01% ± 22.72% | -19.48% ± 4.65% | -32.05% ± 18.16% | Kruskal-Wallis | 0.4038 |
|  | CW | -31.64% ± 0.00% | -24.19% ± 18.87% | -44.63% ± 24.64% | Kruskal-Wallis | 0.2385 |
| Increased | 500 Hz | 76.09% ± 70.44% | 85.28% ± 69.56% | 61.24% ± 42.66% | Kruskal-Wallis | 0.6206 |
|  | CW | 83.07% ± 62.23% | 111.74% ± 112.07% | 58.71% ± 62.87% | Kruskal-Wallis | 0.0966 |

**Supplementary Table S4.** Relative change in firing rates of single units with suppressed or increased activity across cortical layers during pulsed (500 Hz) and continuous-wave (CW) stimulation in wild-type mice (mean ± standard deviation). Statistical comparisons between layers and conditions are reported, including the statistical tests used and corresponding p-values.

|  |  | **BL** | | **ON** | **Statistical test** | **p-value** |
| --- | --- | --- | --- | --- | --- | --- |
| 500 Hz | Theta | | 725.07 μV^2^ ± 238.66 μV^2^ | 700.73 μV^2^ ± 189.43 μV^2^ | Wilcoxon signed-rank | 0.8125 |
|  | Alpha | | 278.31 μV^2^ ± 178.36 μV^2^ | 253.98 μV^2^ ± 132.70 μV^2^ | Wilcoxon signed-rank | 0.8125 |
|  | Beta | | 711.05 μV^2^ ± 500.97 μV^2^ | 562.82 μV^2^ ± 299.14 μV^2^ | Wilcoxon signed-rank | 0.3125 |
|  | Gamma | | 338.71 μV^2^ ± 289.48 μV^2^ | 266.93 μV^2^ ± 181.91 μV^2^ | Wilcoxon signed-rank | 0.8125 |
| CW | Theta | | 695.83 μV^2^ ± 288.56 μV^2^ | 704.41 μV^2^ ± 238.24 μV^2^ | Wilcoxon signed-rank | 0.6250 |
|  | Alpha | | 260.03 μV^2^ ± 147.84 μV^2^ | 234.92 μV^2^ ± 123.19 μV^2^ | Wilcoxon signed-rank | 0.3125 |
|  | Beta | | 709.11 μV^2^ ± 460.73 μV^2^ | 548.83 μV^2^ ± 349.77 μV^2^ | Wilcoxon signed-rank | 0.1250 |
|  | Gamma | | 291.75 μV^2^ ± 189.97 μV^2^ | 241.14 μV^2^ ± 169 80 μV^2^ | Wilcoxon signed-rank | 0.3125 |

**Supplementary Table S5.** Spectral power of theta, alpha, beta, and gamma activity during baseline (BL) and 500 Hz and continuous-wave (CW) stimulation (ON) in wild-type mice (mean ± standard deviation). Statistical comparisons between conditions are reported, including the statistical tests used and corresponding p-values.

|  | **Narrow-spiking interneurons [%]** | **Wide-spiking interneurons [%]** | **Principal cells**  **[%]** |
| --- | --- | --- | --- |
| **KO Mouse 1** | 27.06 | 16.47 | 56.47 |
| **KO Mouse 2** | 37.14 | 20.00 | 42.56 |
| **KO Mouse 3** | 26.26 | 23.23 | 50.51 |
| **KO Mouse 4** | 19.75 | 28.02 | 52.23 |
| **KO Mouse 5** | 21.85 | 24.05 | 53.65 |

**Supplementary Table S6.** Percentages of narrow-spiking interneurons, wide-spiking interneurons, and principal cells identified in each TRPV1 knockout (KO) mouse.

|  |  |  | **WT** | **KO** | **Statistical test** | **p-value** |
| --- | --- | --- | --- | --- | --- | --- |
| Rise time | Suppressed | N-I | 22.00 s ± 11.31 s | 30.33 s ± 11.48 s | Mann-Whitney U | 0.6071 |
|  |  | W-I | 23.00 s ± 4.24 s | 29.50 s ± 15.00 s | Mann-Whitney U | 0.6667 |
|  |  | P | 33.00 s ± 9.89 s | 28.00 s ± 6.00 s | Mann-Whitney U | 0.2793 |
|  | Increased | N-I | 42.64 s ± 11.88 s | 33.58 s ± 13.12 s | Mann-Whitney U | 0.0366 |
|  |  | W-I | 36.86 s ± 13.70 s | 39.56 s ± 15.54 s | Mann-Whitney U | 0.6925 |
|  |  | P | 38.98 s ± 15.34 s | 34.64 s ± 13.25 s | Mann-Whitney U | 0.1860 |
| Fall time | Suppressed | N-I | 18.50 s ± 4.95 s | 15.00 s ± 2.76 s | Mann-Whitney U | 0.2500 |
|  |  | W-I | 24.00 s ± 2.83s | 21.00 s ± 15.44 s | Mann-Whitney U | 0.4667 |
|  |  | P | 21.80 s ± 11.99 s | 19.43 s ± 10.31s | Mann-Whitney U | 0.6833 |
|  | Increased | N-I | 19.93 s ± 10.23 s | 24.61 s ± 19.85 s | Mann-Whitney U | 0.7490 |
|  |  | W-I | 16.00 s ± 7.30 s | 20.86 s ± 11.43 s | Mann-Whitney U | 0.3031 |
|  |  | P | 19.62 s ± 9.92 s | 21.34 s ± 11.17 s | Mann-Whitney U | 0.5209 |

**Supplementary Table S7.** Rise and fall times of narrow-spiking interneurons (N-I), wide-spiking interneurons (W-I), and principal cells (P) with suppressed or increased activity during continuous-wave (CW) stimulation in wild-type (WT) and TRPV1 knockout (KO) mice (mean ± standard deviation). Statistical comparisons between conditions are reported, including the statistical tests used and corresponding p-values.

|  | **Layers 1/2/3/4** | **Layer 5** | **Layer 6** |
| --- | --- | --- | --- |
| **KO Mouse 1** | 7 | 50 | 28 |
| **KO Mouse 2** | 32 | 0 | 3 |
| **KO Mouse 3** | 23 | 54 | 22 |
| **KO Mouse 4** | 18 | 58 | 81 |
| **KO Mouse 5** | 17 | 50 | 84 |
| **Total** | 97 | 212 | 218 |

**Supplementary Table S8.** Number of single units detected in each cortical layer group for each TRPV1 knockout (KO) mouse.

|  |  | **WT** | **KO** | **Statistical test** | **p-value** |
| --- | --- | --- | --- | --- | --- |
| Layers  1-4 | Suppressed | 1.18% ± 2.63% | 6.72% ± 9.48% | Mann-Whitney U | 0.4444 |
|  | Increased | 29.11% ± 31.63% | 40.00% ± 22.92% | Mann-Whitney U | 0.7381 |
| Layer 5 | Suppressed | 27.04% ± 41.90% | 2.93% ± 4.55% | Mann-Whitney U | 0.2857 |
|  | Increased | 26.99% ± 32.29% | 44.82% ± 30.59% | Mann-Whitney U | 0.4127 |
| Layer 6 | Suppressed | 5.86% ± 11.68% | 5.43% ± 7.43% | Mann-Whitney U | >0.9999 |
|  | Increased | 32.08% ± 11.75% | 34.05% ± 23.85% | Mann-Whitney U | 0.8889 |

**Supplementary Table S9.** Percentage of single units detected in each cortical layer group with suppressed or increased activity in wild-type (WT) and TRPV1 knockout (KO) mice (mean ± standard deviation). Statistical comparisons between conditions are reported, including the statistical tests used and corresponding p-values.

|  | **WT** | | **KO** | **Statistical test** | **p-value** |
| --- | --- | --- | --- | --- | --- |
| Delta | | 28.14% ± 12.60% | 49.86% ± 33.83% | Mann-Whitney U | 0.3095 |
| Theta | | 12.73% ± 11.91% | 28.14% ± 15.68% | Mann-Whitney U | 0.1508 |
| Alpha | | -6.33% ± 13.31% | 0.38% ± 22.09% | Mann-Whitney U | 0.6905 |
| Beta | | -13.84% ± 9.50% | -12.36% ± 5.52% | Mann-Whitney U | 0.6905 |
| Gamma | | 6.26% ± 5.57% | -2.93% ± 11.03% | Mann-Whitney U | 0.3095 |

**Supplementary Table S10.** Change in the power of delta, theta, alpha, beta, and gamma bands during continuous-wave (CW) stimulation in wild-type (WT) and TRPV1 knockout (KO) mice relative to baseline (mean ± standard deviation). Statistical comparisons between WT and KO animals are reported, including the statistical tests used and corresponding p-values.
